# Claustral Projections to Anterior Cingulate Cortex Modulate Engagement with the External World

**DOI:** 10.1101/2021.06.17.448649

**Authors:** Gal Atlan, Noa Matosevich, Noa Peretz-Rivlin, Idit Yvgi, Eden Chen, Timna Kleinman, Noa Bleistein, Efrat Sheinbach, Maya Groysman, Yuval Nir, Ami Citri

## Abstract

Engagement is a major determinant of performance. Hyper-engagement risks impulsivity and is fatiguing over time, while hypo-engagement could lead to missed opportunities. Even in sleep, when engagement levels are minimal, sensory responsiveness varies. Thus, maintaining an optimal engagement level with the environment is a fundamental cognitive ability. The claustrum, and in particular its reciprocal connectivity with executive regions in the frontal cortex, has been associated with salience, attention and sleep. These apparently disparate roles can be consolidated within the context of engagement. Here we describe the activity of claustro-frontal circuits in a task imposing a tradeoff between response inhibition and sensory acuity (‘ENGAGE’). Recording calcium fiber photometry during >80,000 trials, we characterize claustrum recruitment during salient behavioral events, and find that a moderate level of activity in claustro-cingulate projections defines optimal engagement. Low activity of this pathway is associated with impulsive actions, while high activity is associated with behavioral lapses. Chemogenetic activation of cingulate-projecting claustrum neurons suppressed impulsive behavior and reduced the engagement of mice in the task. This relationship became even clearer upon addressing individual variability in the strategy mice employed during the ENGAGE task. Furthermore, this association of claustrum activity and engagement extends into sleep. Using simultaneous EEG and photometry recordings in the claustrum, we find that cingulate projecting claustrum neurons are most active during deep unresponsive slow-wave sleep, when mice are less prone to awakening by sensory stimuli.

## Introduction

Engagement is a crucial determinant of behavior. Sensory events that are normally ignored can become highly salient, depending on attentional state and engagement with the external world. In the 2011 World Championships, reigning champion and world record holder Usain Bolt committed a false start on the 100 meter final and was disqualified from the competition. It cannot be said that Bolt lacked experience or skill, as he is widely regarded as the best sprinter of all time ^1^. In fact, his reaction time is considered slow, an indication of his confidence and natural sprinting ability ^2^. As he was waiting for the gun to start the race, the slightest twitch of a muscle from his compatriot and eventual race winner, Yohan Blake, was arguably the trigger that sent him off prematurely. Bolt’s hyper-sensitivity likely reflects his vigilant concentration and anticipation of the start signal, amplified by the high-pressure occasion.

In challenging tasks or under pressure, performance scales with engagement only up to a limit, following a bell-shaped curve known as the ‘Yerkes Dodson’ law ^3^. Optimal performance is achieved when a balance between vigilance and caution is achieved (being “in the zone”). Hyper-engagement reduces task performance by increasing the propensity for impulse errors, while hypo-engagement (“zoning out”) leads to missed opportunities. Furthermore, maintenance of heightened engagement over time eventually leads to exhaustion and lapses in attention ^4,5^. Engagement with the external world can also be addressed in sleep, as deeper sleep is associated with a reduced propensity to be awoken by sensory stimuli ^6,7^.

Prefrontal regions of the cortex, such as the anterior cingulate cortex (ACC), and the orbitofrontal cortex (OFC), have been implicated in regulating multiple attentional processes such as vigilance and impulse control, positioning them as prime candidates for modulating engagement with the external world ^8-11^. Prefrontal cortex is heavily modulated by global arousal signals, widely attributed to the action of neuromodulators and to subcortical structures such as the thalamus and the claustrum, due to their capacity to synchronously signal to broad cortical territories ^12-15^. In this study, we identify a role for claustral neurons projecting to the ACC in defining the degree to which mice engage with the external world.

The claustrum is a thin neuronal structure, enclosed between the insular cortex and the striatum in the mammalian brain ^15-17^. It has been proposed to mediate cortical synchronicity, salience and attention ^15,18-22^ through strong claustro-cortical feed-forward inhibition ^14,19,23^. The most prominent connectivity of the claustrum is with prefrontal cortical structures such as the ACC and OFC ^24-28^. Axons from these frontal regions reciprocally innervate most of the claustrum, in contrast to constrained sensory zones defined by afferents from sensory cortices ^26,27,29-33^. Further anatomical division of the claustrum into modules is supported by its internal organization into a ‘core’ and ‘shell’, as well as by mapping of its projections ^24,25,32,34,35^. Such modules could potentially play distinct roles in modulating executive function and sensory processing, particularly given recent studies associating claustrum activity with behavioral performance ^19,20,36^. However, the association between claustral modules and physiology and function is yet to be clearly demonstrated ^37^. Particularly, data regarding the response patterns of claustral populations during behavior are scarce ^18,36,38^, and the rules governing behaviorally-relevant claustrum recruitment remain unexplored.

Here we employed fiber photometry from anatomically defined claustral projection networks, recording calcium transients in behaving mice. Our results demonstrate that ACC-projecting (ACCp) and OFC-projecting (OFCp) claustrum neurons form distinct modules, differing in their anatomical distribution as well as in their spontaneous activity and their recruitment during behavior. Utilizing an automated behavioral training system, we trained mice on a cognitively-engaging task (‘ENGAGE’), imposing a tradeoff between response inhibition and engagement. We find that claustro-frontal populations were recruited during the task, responding transiently to multiple salient sensory events and motor actions. Importantly, the activity of ACCp neurons, but not OFCp neurons, reflects the level to which mice engage with the task. Thus, low ACCp activity corresponds to hyper-engagement, while high ACCp corresponds to disengagement. Chemogenetic elevation of the activity of ACCp neurons was sufficient to suppress impulsive responses. Furthermore, we observed that mice exhibited distinct strategies for coping with the ENGAGE task, which related to their degree of ACCp recruitment. Finally, by applying simultaneous EEG and photometry recordings, we found that the association between ACCp activity and engagement extends to sleep. Claustrum activity increased during periods of maximal slow wave activity (SWA) in NREM sleep, and correlated with the potential of a mouse to maintain its sleep in the presence of awakening tones. Taken together, our results reveal the role of a sub-network of claustral neurons projecting to the ACC in controlling engagement with the external world (Figure S1).

## Results

### Claustral subpopulations projecting to ACC vs OFC are largely distinct

To investigate the functional organization of claustrum subpopulations projecting to frontal regions, we focused on two main frontal targets of the claustrum: the ACC and the OFC. Projection-based labeling of claustral neurons, enabled by retrograde-transporting Adeno Associated Viruses^39^ (retroAAV; Figure 1A, and see supplementary table T1), demonstrated that ACCp and OFCp claustrum neurons are anatomically segregated (Figure 1B, S2A), with sparse co-labeled neurons (Figure S2B). ACCp neurons are densely clustered in the claustrum core, while OFCp neurons are spread throughout the core and the shell of the claustrum (Figure 1C, S2C-D), appearing sparser in the core (Figure S2A, D). Axons of ACCp and OFCp claustral neurons exhibit differential projections to frontal targets (Figure 1D). In addition, ACCp axonal arborization was more prominent within sensory cortical areas (Figure 1E, S2E).

**Figure 1.**
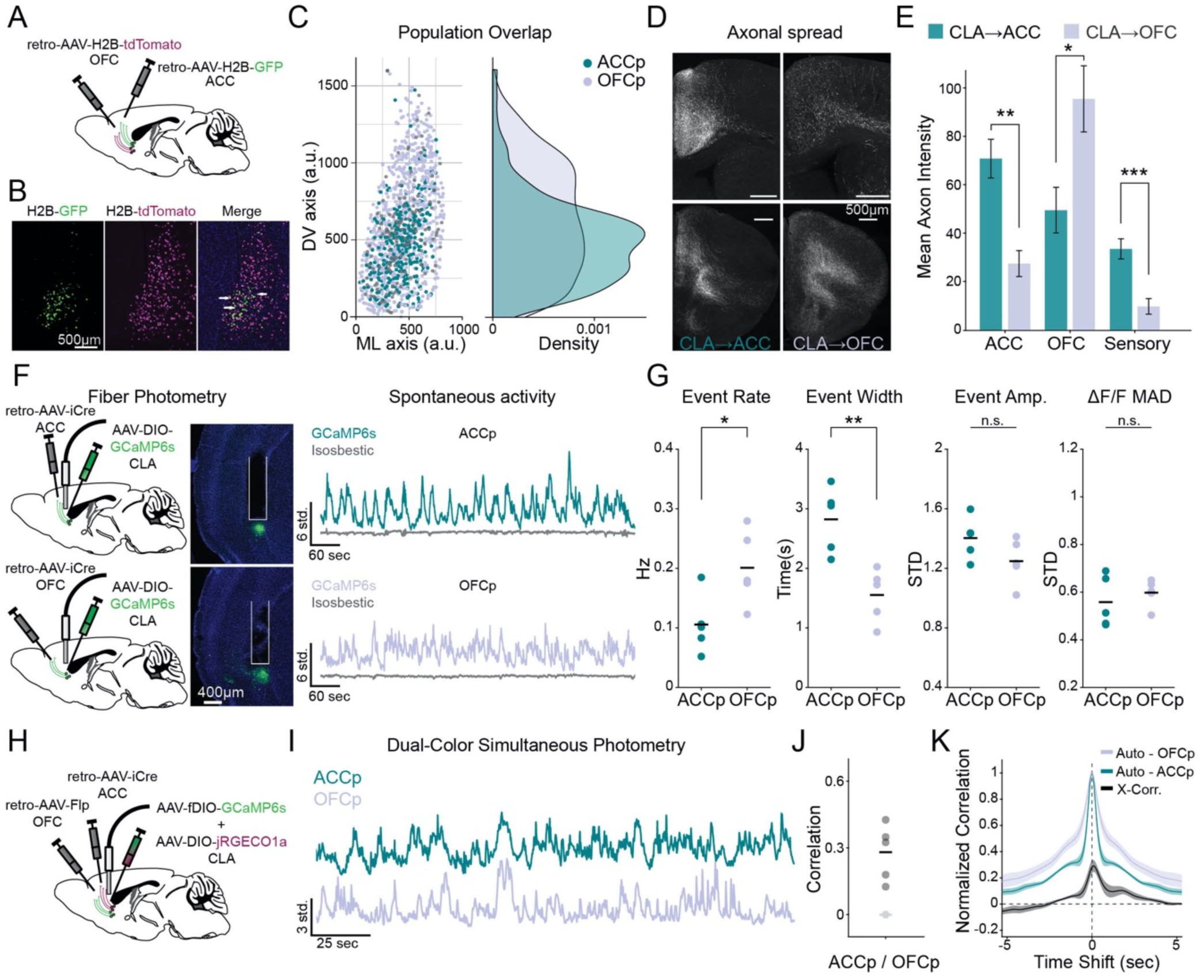
Differential claustrum networks project to ACC vs OFC. (A) Scheme for dual-color soma-targeted retrograde labelling of claustrum projection neurons. (B) Example expression of H2B-GFP in ACC-projecting neurons (ACCp; left); H2B-tdTomato in OFC-projecting neurons (OFCp; middle); and double-labeled neurons (right; white arrows). (C) Digitized overlap of all neurons from a single coronal plane (+0.38mm relative to Bregma) over all mice (n=3), and their distribution of expression along the dorsoventral axis of the claustrum (right). Dark gray indicates double-labeled cells. (D) IHC-amplified GFP-labelled ACCp (left) or OFCp (right) axonal projections within ACC (top) and OFC (bottom). See (F) for viral approach. (E) Mean fluorescence intensity in ACCp (n=4 mice) and OFCp (n=3 mice) projections. (F) Approach for fiber photometry recordings from ACCp (top) vs OFCp (bottom) claustrum populations. Middle panels depict representative histological expression and optic fiber placement. Right panels depict spontaneous activity in head-restrained mice. (G) Quantification of spontaneous calcium event rate, width (at half maximal prominence), amplitude, and overall median absolute deviation (MAD) of ACCp vs OFCp (n=5 mice in each group) z-scored ΔF/F. (H) Approach for simultaneous recording from ACCp and OFCp neurons using two-color photometry. (I) Representative spontaneous photometry traces from an ACCp/OFCp mouse. (J) Correlation between spontaneous co-activity in ACCp/OFCp mice (n=5). Light gray dot represents the maximal correlation over 1000 iterations of shuffled data per mouse, averaged across mice. (K) Average cross-correlation of spontaneous activity in ACCp/OFCp mice (gray, n=5) in comparison to the auto-correlations of OFCp (purple) and ACCp (turquoise). Unless noted otherwise, data are mean ± s.e.m. *p<0.05, **p<0.01, ***p< 0.001; n.s., not significant. See Supplementary Table 3 for further details of statistical analyses.

To address whether the anatomical segregation between ACCp and OFCp neurons extends also to their physiology, we employed an intersectional viral approach to record population calcium transients from ACCp or OFCp neurons in head-restrained mice running on a linear track (Figure 1F; Methods). ACCp activity exhibited less frequent but longer-lasting spontaneous calcium events in comparison to OFCp (Figure 1G). We next recorded concurrently from both populations in the same animal, using dual-color fiber photometry (Figure 1H, I). Spontaneous activity was correlated between ACCp and OFCp neurons (Figure 1J). However, these windows of correlation were relatively short, as cross-correlations of ACCp and OFCp activity decayed within a second (Figure 1K). Together, these results establish the ACCp and OFCp subpopulations as partially overlapping, yet largely independent claustro-frontal networks.

### ACCp activity bi-directionally reflects task engagement

We next proceeded to investigate the recruitment and function of the ACCp and OFCp populations during behavior. We developed ‘ENGAGE’, a novel biphasic form of a randomized cue delay task, supporting the investigation of multiple aspects of attentive behavior, including impulsivity, sensory detection and selection, and sustained attention ^40^. Trial onset (initiated by the mouse during training, or every 20s during recording sessions, see below) was indicated by a brief broadband noise (BBN), followed by a randomized delay period (of 0.5-3s), during which mice were required to withhold licking until an auditory ‘go’ cue was played. Premature (‘impulsive’) licking during this initial stage of the task resulted in trial abortion. Timely licks (<1.5s) following the go-cue were rewarded (‘hit’). Trials in which the mouse did not lick in a timely fashion were defined as ‘miss’ trials and were not rewarded (Figure 2A; Methods). Trial difficulty was determined by a combination of several conditions: go-cue tone (four intensities), a tone cloud distractor (presence, absence), and a visual aid, presented together with the auditory go-cue (presence, absence). Conditions were randomized across trials, while maintaining fixed proportions of each condition throughout sessions (Figure 2B). Mice were trained in a custom automated behavioral setup, supporting simultaneous individualized training of multiple mice in their home cage (Figure S3 and see Methods). Upon completion of the automated training protocol in their home cage, mice transitioned to a head-restrained setup for individual recordings, allowing for well-controlled photometry recordings over numerous trials (∼100,000 trials from 30 mice in sum, see supplementary table T1). Mice reliably transferred their learning from the automated training to the head-fixed condition (Figure 2C). The ENGAGE task was designed to probe fluctuations of hyper- and hypo-engagement, supported by high proportions of both impulsive and omission errors (Figures 2D). On average, the performance of mice was impacted by all trial variables: improving psychometrically as a function of increased intensity of the auditory cue, while performance at low cue intensities benefitted from the addition of the visual aid. The tone cloud contributed to overall attentional load, resulting in reduced hit rates, primarily in trials with intermediate cue intensities, as well as directly increasing impulsive error rates (Figure 2E).

**Figure 2.**
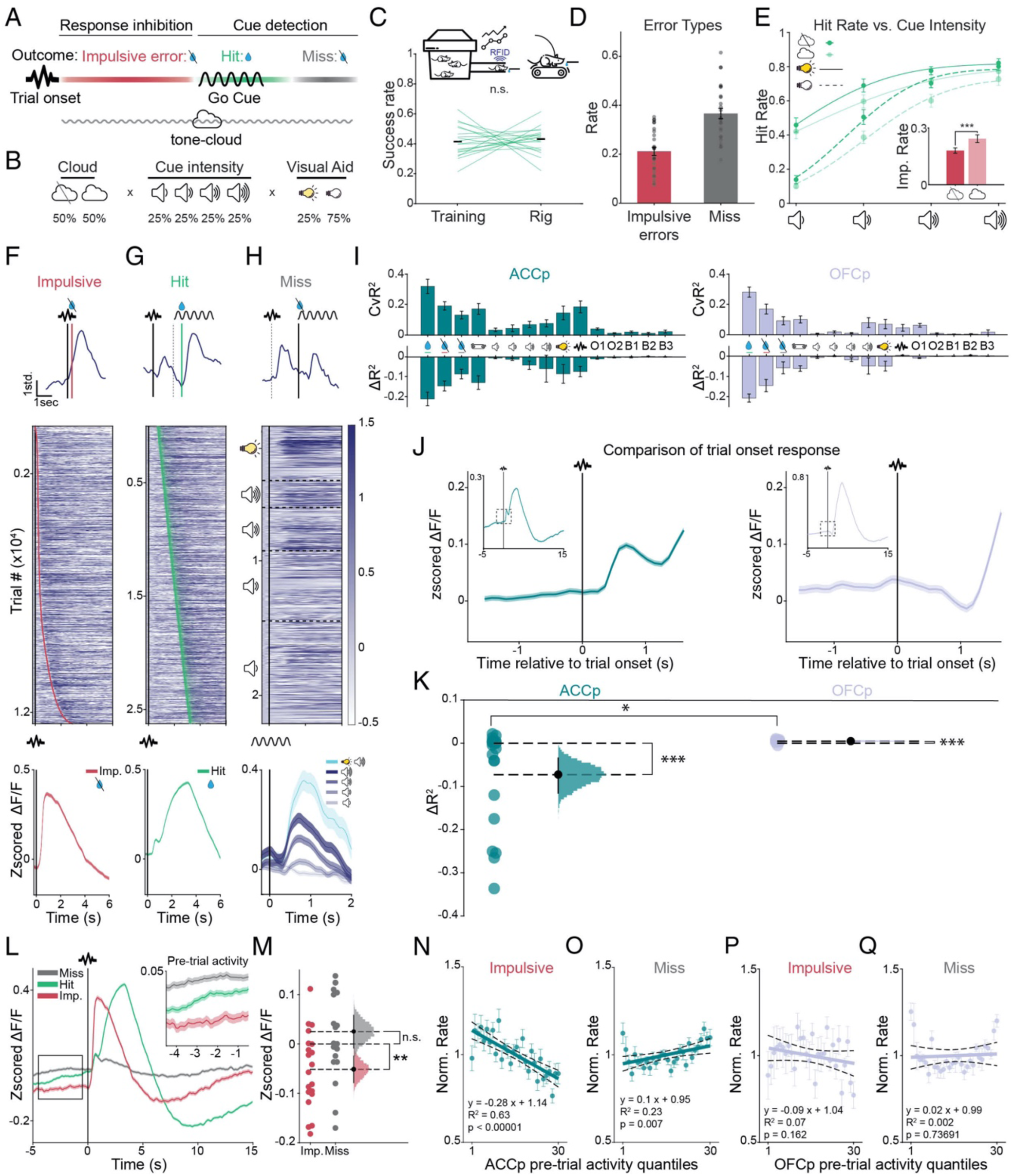
Differential correlation of ACCp vs OFCp activity with trial outcome. (A) Scheme describing the ENGAGE task. Trial onset was indicated by a 100ms broadband noise. Mice were rewarded for timely responses following Go cue initiation (‘hit’). Impulse errors were defined as licks between trial onset to the Go cue, while ‘miss’ trials included trial omissions and late licks. (B) Distribution of trial parameters. (C) Co-housed cohorts of mice (n=23 mice in 7 cages) were trained in an automated home-cage system (see Methods), allowing individualized training schemes based on RFID identification. Success rate transferred from training to subsequent head-fixed recording sessions. (D) Distribution of impulsive vs miss error rates in the task during head-fixed recording sessions. (E) Mean hit rate (excluding impulsive errors) as a function of cue intensity during recording (∼80,000 total individual trials). Inset depicts impulse errors, which increased in the presence of the tone cloud. (F-H) ACCp claustrum dynamics during impulsive (F, n=12,471), hit (G, n=26,243) and miss (H, n=23,430) trials aligned to trial onset (F-G) or cue presentation (H). Top: Single trial examples. Red and green lines indicate impulsive or correct licks, respectively. Heatmaps: all ACCp trials from n=20 mice, sorted by lick onset (impulsive); the delay from trial onset to cue (hits); or cue intensity (miss). Ticks indicate the first impulsive or correct lick within the trial, respectively. Bottom: mean activity traces in impulsive (left) hit (middle) and miss trials (right, separated by cue intensity). (I) Quantification of the contribution of behavioral events to claustrum photometry signal (n=20 ACCp channels, n=10 OFCp channels) using a linear encoding model (see Methods and supplementary table T2). CvR^2^: cross-validated explained variance in a single variable model compared to the full model. ΔR^2^: unique contribution of a label to the model measured by net loss in explained variance. (J) Averaged ACCp (left, n=20 channels) and OFCp (right, n=10 channels) traces around trial onset. (K) Model quantification of the representation of trial onset. Data is presented as individual mice with bootstrapped distribution of means, and 95% confidence intervals. (L) Mean activity of all ACCp recordings (n=20) aligned to trial onset, separated by trial outcome. Inset depicts pre-trial activity. (M) Mean pre-trial activity preceding impulsive (red) or miss (gray) errors (individual mice with bootstrapped distribution of means and 95% confidence intervals). (N-Q) Normalized impulsive (N,P) or miss (O,Q) error rate, as a function of pre-trial activity quantiles for AACp (N,O; n=20) or OFCp (P,Q; n=10) data. Thick line represents linear fit, dotted lines represent 95% confidence intervals. Unless noted otherwise, data are mean ± s.e.m. *p<0.05, **p<0.01, ***p< 0.001; n.s., not significant. See Supplementary Table 3 for further details of the statistical analyses.

Within the ENGAGE task, both ACCp and OFCp claustrum populations were recruited by impulsive as well as correct licks (Figures 2F-G, S4A, B). In contrast, ‘miss’ trials provided a window into the claustral representation of the go-cue in the absence of confounding lick events (Figures 2H, S4C). In order to quantify the degree to which discrete temporal epochs contributed to the activity of claustral networks, we fit a linear encoding model to the data ^41^, creating a time-varying event kernel to relate each event to its corresponding neural signal. We then compared the cross-validated explained variance (CvR^2^) for each event independently, as well as the unique contribution (AR^2^) of that event to explaining the total calcium signal (Figure 2I, S5A, B; supplementary table T2 and Methods). This analysis served to quantify claustral activity with relation to particular behavioral events, revealing that, as reported in the cortex ^41,42^, claustrum activity in both ACCp and OFCp networks was evoked by spontaneous locomotion (Figure S5C); task-related licking (Figure S5D); and, in some mice, by sensory stimuli (Figure S5E). Strikingly, trial onset was strongly represented in ACCp, but not in OFCp activity (Figure 2J, K). Indeed, trial onset appears to have been a significant catalyst of impulsivity, as 82% of impulsive errors occurred within 1 second of the BBN signaling trial onset (Figure 2F). The unique coupling of the ACCp signal to this major determinant of impulse errors suggested that that the ACCp network may function to modulate response inhibition.

We therefore wished to determine whether ACCp activity leading up to trial onset may vary in preparation for this predictable, yet challenging aspect of trial structure. Plotting ACCp activity by trial outcome (Figure 2L), we observed that pre-trial ACCp activity was lower on average preceding trials terminated by impulsive errors, compared to hit trials (Figure 2M). In fact, pre-trial ACCp activity exhibited an inverse correlation with impulsive errors, such that lower pre-trial ACCp activity corresponded to a higher probability that the trial would result in a premature lick (Figure 2N). Intriguingly, the opposite relationship was observed between pre-trial ACCp activity and miss errors, such that trials with higher pre-trial ACCp activity were more likely to result in misses (Figure 2O). OFCp pre-trial activity was not significantly lower before impulsive errors (Figure S6A, B), nor was it correlated with impulse errors (Figure 2P) or misses (Figure 2Q). Importantly, we observed no correlation between pre-trial ACCp activity and reaction time in hit trials, suggesting that ACCp activity related specifically to the capacity of mice to engage with the trial, rather than correlating with disruptions to perception or action (Figure S6C, D). In sum, while ACCp and OFCp claustral networks are recruited during multiple stages of the task, ACCp activity was uniquely tied to trial onset and to the propensity of mice to engage in impulsive licking following low activity levels, or misses following high activity.

### Elevating ACCp activity reduces impulsive action

In order to address the causal role ACCp activity plays in controlling impulsive behavior and engagement, we co-expressed the excitatory DREADD hM3Dq together with GCaMP6s in ACCp neurons (n=5, see supplementary table T1). This enabled a direct measurement of the effects of chemogenetic manipulation on ACCp activity within the context of the task (Figures 3A, S7A). Mice underwent behavioral training as described above, and were habituated to saline injections during head-restrained behavioral sessions. CNO administration (10mg/kg, i.p.) reliably elevated spontaneous claustrum activity (Figure 3B-C). Mice were then tested following administration of either saline or CNO on interleaved days. Strikingly, CNO significantly and reversibly reduced impulsive error rates, implicating the ACCp in control of impulsivity (Figure 3D). CNO administration did not change impulsive error rates in GCAMP6s controls (Figure S7B). It lead to no changes in the representation of task parameters in the ACCp signal in hM3Dq mice (Figure S7C), nor did CNO affect the response times of mice in hit trials or their overall success rate (Figure S7D, E). Intriguingly, CNO administration induced a shift in the distribution of trial outcomes over the course of a session, such that in the first half of the session, impulsive errors were largely replaced by hits, while in the second half of the session, miss trials were more common (Figure 3E). Consistent with interpreting an elevation of miss trials as a decrease in engagement, streaks of consecutive miss trials were more common following CNO (Figure S7F). Thus, chemogenetic induction of ACCp activity reduced impulsivity at the cost of increased miss trials over prolonged sessions.

**Figure 3.**
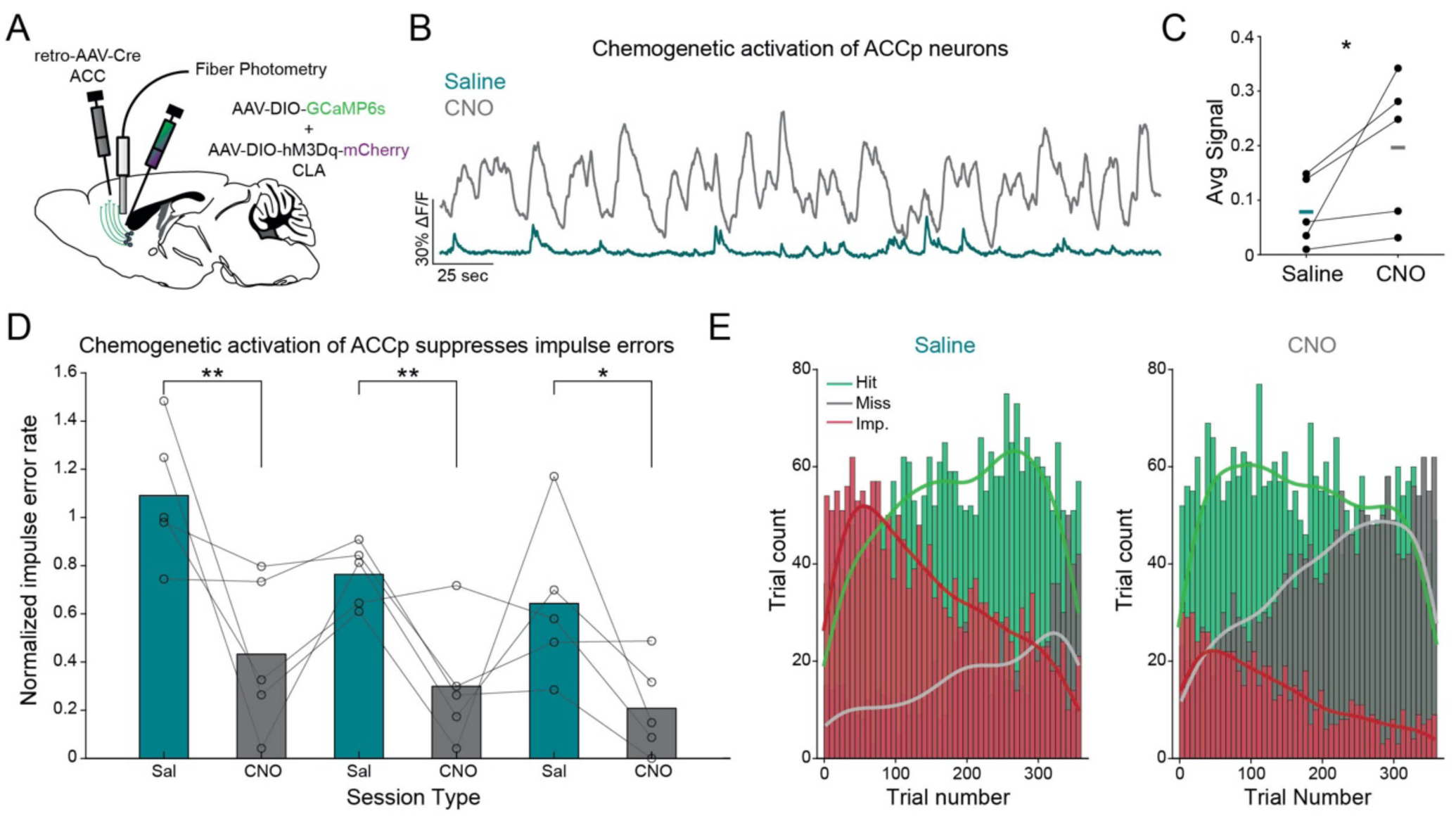
Chemogenetic facilitation of ACCp activity diminishes impulsive behavior. (A) Approach for simultaneous chemogenetic activation and recording of ACCp activity. (B) Example recordings of spontaneous activity from an ACCp mouse following saline (turquoise) or CNO (gray) administration. (C) Average spontaneous calcium signal following saline vs CNO (10mg/kg i.p) administration (n=5 mice). (D) Comparison of impulsive errors in interleaved daily sessions of saline vs CNO, normalized to the average rate over 3 prior days of saline habituation (n=5 mice). (E) Binned histograms (vertical lines) and kernel fit (smooth horizontal lines) of the distribution of trial outcome within saline (left) vs CNO (right) sessions (n=3 sessions/each from 5 mice; 360 trials/session). Unless noted otherwise, data are mean ± s.e.m. *p<0.05, **p<0.01, ***p<0.001; n.s., not significant. See Supplementary Table 3 for further details of the statistical analyses.

### ACCp activity reflects attentional strategy

We next addressed the distribution of behavioral strategies taken by mice in dealing with the ENGAGE task. We divided mice into three behavioral categories based on the degree to which their behavior was affected by task parameters (Figure 4A, B). 5/25 mice exhibited a selective approach, primarily participating in easy trials with prominent cues or a visual aid *(selective;* cue modulation index >0.5). A second group of mice (6/25) exhibited behavior that was consistent across trial parameters *(consistent;* cue modulation index <0.5), and scaled with cue intensity. Both groups were minimally affected by the cloud distractor (cloud modulation index < 0.04). In contrast, the largest group of mice (14/25), was more susceptible to interference by the cloud *(erratic;* cloud modulation index >0.04). We next addressed whether these groups corresponded to other elements of behavior in the task. *Consistent* mice exhibited the highest overall success rate (mean rate 51%), distinguished from *erratic* mice, whose success rate was lowest (mean rate 38%) (Figure S8A). The groups also differed in their impulsive error rates, with *erratic* mice exhibiting a higher probability of performing impulsive licks, which further increased in the presence of the tone cloud (Figure 4C). As noted earlier, impulsive errors closely coupled to trial onset. This effect varied with the strategy of mice, and was most prominent in *erratic* mice, evident in faster response times in impulsive errors (Figure 4D). Importantly, response times in hit trials did not differ between groups, suggesting that potential confounds, relating to perception or motor deficiencies across behavioral categories, are unlikely (Figure S8B). Thus, *erratic* mice appeared to be hyper-engaged with the task, exposing them to impulsive erroneous responses to the trial-onset cue and the cloud.

**Figure 4.**
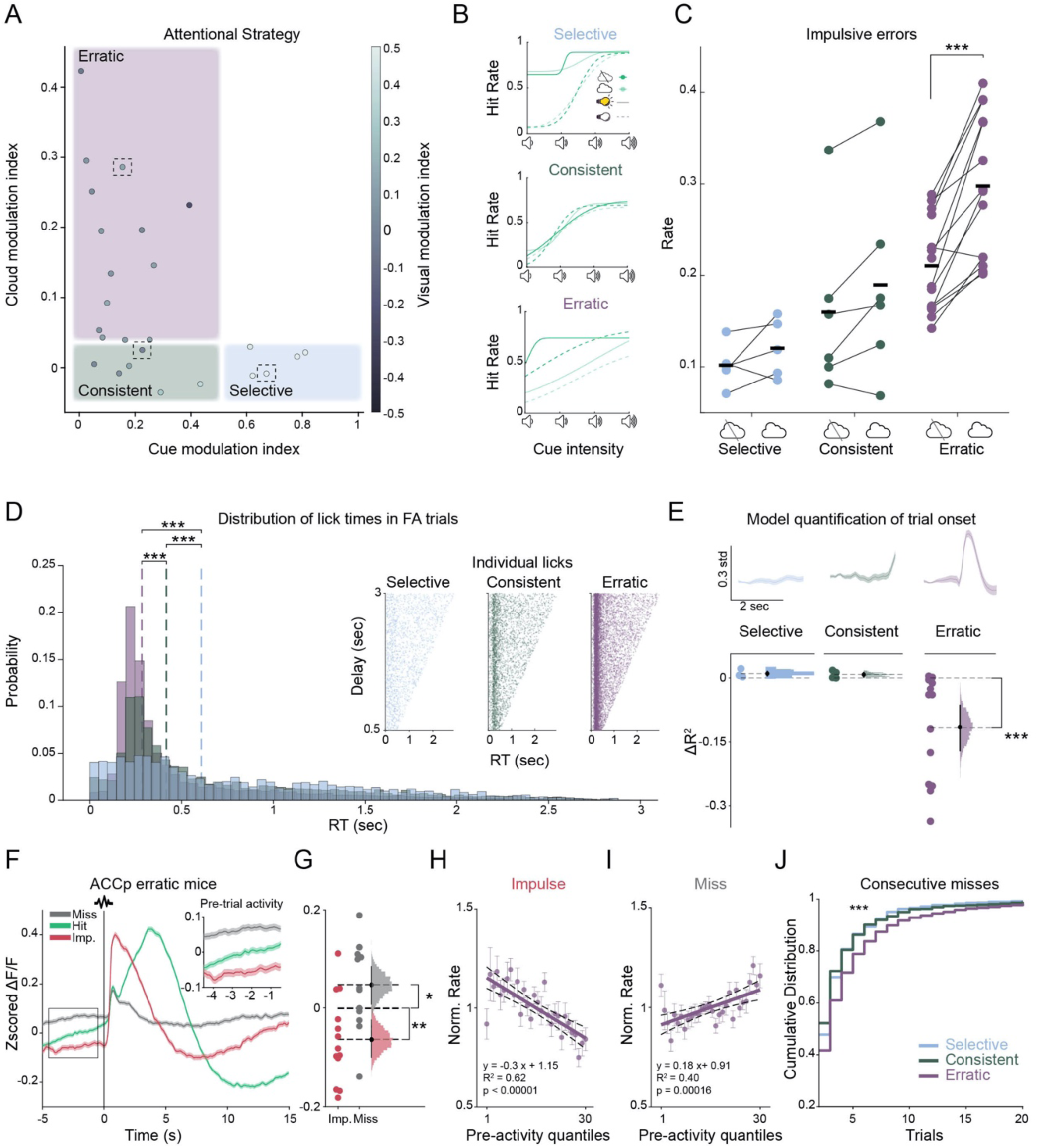
ACCp activity corresponds to individual differences in behavioral strategies. (A) Individual mice are plotted according to their modulation indices depicting the dependency of individual hit rates on cue intensity (cue modulation index), cloud (cloud modulation index), or visual aid (visual modulation index, represented by the shading of the dots). Mice (n=25) were grouped into three groups, based on their strategy in the task (‘selective’= cue modulation index>0.5; ‘consistent’= cloud modulation index<0.04 & ‘erratic’; n=5, 6, 14, respectively). (B) Psychometric curves of representative mice from each group (dotted frames in A). (C) Impulse error rate in absence or presence of the tone-cloud, by behavioral group. (D) Distribution of impulsive lick response times. Dotted lines indicate distribution medians. Inset depicts all trials as a function of the random delay period. (E) Representation of trial onset in the ACCp signal. Top: Average responses in representative ACCp signals. Bottom: Model quantification of trial onset response. Individual mice (n=3, 4, 13) and bootstrapped distribution of means with 95% confidence intervals. (F) Mean activity in ACCp recordings from erratic mice (n=13) aligned to trial onset, separated by outcome. Inset depicts pre-trial activity. (G) In *erratic* mice the ACCp activity preceding impulsive errors is low, while ACCp activity preceding miss trials is high. Individual mice and bootstrapped distribution of means with 95% confidence intervals. (H-I) Normalized impulse (H) or miss (I) error rate, as a function of pre-trial activity of ACCp in erratic mice. Thick line represents linear fit, dotted lines represent 95% confidence intervals. (J) Cumulative distribution of consecutive miss trials for each behavioral group. Unless noted otherwise, data are mean ± s.e.m. *p<0.05, **p<0.01, ***p< 0.001; n.s., not significant. See Supplementary Table 3 for further details of the statistical analyses.

Consistent with the coupling of impulsive errors and ACCp activity, we observed an inverse correlation between the response time in impulsive trials and the unique contribution of trial onset to the ACCp signal (Figure S8C). Furthermore, *erratic* mice, which were the most prone to impulsive errors, showed the strongest ACCp response to trial onset (Figure 4E). In light of this, we re-addressed ACCp pre-trial activity, specifically in *erratic* mice. The association between trial outcome to ACCp pre-trial activity was even more pronounced in this group in comparison to all mice (Figure 4F, G). Maintaining a strong negative correlation between pre-trial activity and impulsivity (Figure 4H), the positive correlation of ACCp pre-trial activity in *erratic* mice with misses was stronger compared to all mice (Figure 4I). *Erratic* mice also made more streaks of consecutive misses compared to the other groups (Figure 4J), consistent with the notion that these mice transition between states of extreme engagement, characterized by low ACCp activity, and periods of ‘zoning out’, characterized by high ACCp activity. In addition, these streaks (>5 consecutive misses) were preceded by an increased reaction time in hit trials (Figure S8D), as may expected by a gradual decrease in engagement (‘zoning out’). Thus, by considering individual differences in strategy within the ENGAGE task, we highlight the bidirectional relationship between ACCp activity and hyper-engagement vs. disengagement.

### Claustrum activity fluctuates on ultra-slow scales, together with inputs from auditory cortex

To understand whether ACCp activity is driven by cortical projections to the claustrum, we performed recordings of axon-targeted GCAMP6s, expressed in inputs to the claustrum from the ACC (ACCi) and auditory cortex (AUDi), together with jRGECO1a activity in ACCp neurons (Figure S9A). Spontaneous correlations of ACCi or AUDi with ACCp were low, suggesting that neither of these inputs is the main driver of spontaneous ACCp activity (Figure S9B). During the ENGAGE task, however, a strong correlation emerged between AUDi and ACCp (0.6±0.1, in comparison to spontaneous correlation of 0.22±0.1). This contrasted with the correlations between ACCp and both OFCp and ACCi, which were maintained at a similar level in the task as during spontaneous recordings (Figure S9C-D). The AUDi signal also represented task events similarly to the ACCp with respect to licking (Figure S9E), cue responses (Figure S9F), and representation of trial onset (Figure S9G). However these transient events are unlikely to account for the increase in correlation between ACCp and AUDi as the cross-correlation between the two signals was maintained over prolonged time (Figure S9C). In fact, pre-trial activity was correlated between the ACCp and AUDi, while no correlation of pre-trial activity was evident between the ACCp and either the OFCp or ACCi (Figure S9H, I). These results suggest that the ACCp and AUDi acquired a common source of slow modulation within the context of the ENGAGE task, from which OFCp and ACCi were exempt. Behavioral sessions lasted up to three hours, enabling analyses of slow periodicity of pre-trial activity during continuous task performance. Indeed, we observed that activity of all recorded channels tended to fluctuate at an ultra-slow time scale, on the order of tens of minutes (0.1-0.7 mHz; Figure S9J). However, not all signals from all mice exhibited these fluctuations, and interestingly, the mice whose ACCp activity lacked ultra-slow fluctuations were associated with the *erratic* group, which employed the least moderated behavioral strategy (Figure S9K). In sum, a strong correlation emerges between the ACCp and AUDi within the context of the ENGAGE task, maintained across ultra-slow time scales, potentially corresponding to a moderated approach to task performance.

### High ACCp activity corresponds to higher slow-wave activity and deeper, unresponsive sleep

Sleep can be considered as the extreme end of the vigilance-disengagement spectrum. Claustroforebrain activity has recently been associated with sleep, particularly with the occurrence of slow wave activity (SWA, EEG spectral power below 4Hz) in mice ^14^. To further investigate how specific claustro-frontal subnetworks are recruited during extreme states of disengagement, we performed simultaneous polysomnography recordings (EEG, EMG, and video) together with fiber photometry from ACCp and OFCp networks (Figure 5A; ACCp n=6, OFCp n=6, Methods, and see supplementary table T1). During daytime ‘lights on’ periods, mice spent 34.4±5.8%, 54±7.1%, and 8.1±1% of their time in wakefulness, NREM sleep, and REM sleep, respectively (Figure 5B-D), in agreement with the literature ^43^. We found that ACCp activity was lowest in REM sleep and highest during NREM sleep, while intermediate activity was observed in wakefulness (Figure 5E). OFCp similarly demonstrated low activity in REM and high activity in NREM. Yet unlike ACCp, activity in this network during wake trended towards even higher levels than those observed during NREM (Figure S10A). We proceeded to examine whether ACCp activity correlates with specific EEG patterns by dividing ACCp activation into quartiles within each state and examining the corresponding EEG power spectrum (Figure 5F; Methods). We observed that different levels of ACCp activity were associated with different profiles of SWA (< 4Hz) and theta frequencies (6 -9Hz). SWA is an established marker of sleep depth ^44^, and theta activity is maximal during active exploration ^45^. Thus, we used the ratio between SWA and theta power as an EEG index for disengagement, and assessed its relation with ACCp activity. We found that ACCp activation exhibited a positive linear relationship with SWA-to-theta ratio in each behavioral state (Figure 5F, see also supplementary table T3 and Methods).

**Figure 5.**
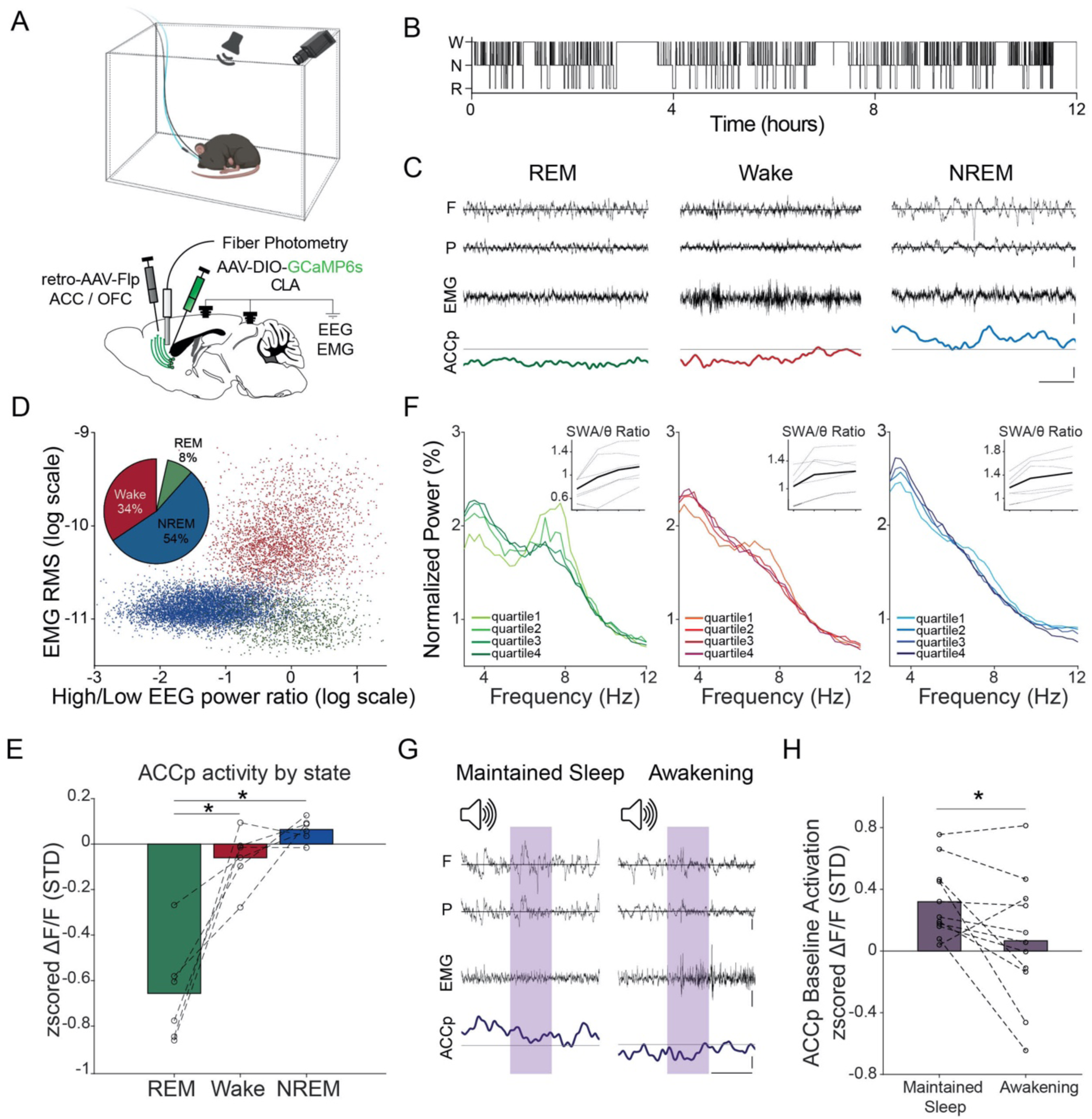
Claustral ACCp activity is tied to deeper NREM sleep. (A) Top: Diagram of experimental setup for recording from a freely behaving mouse in its home-cage under video surveillance, in the presence of a speaker for tone presentations. Bottom: Simultaneous monitoring of frontal and parietal EEG, neck EMG, and fiber photometry from ACCp or OFCp claustral neurons. (B) Representative hypnogram (time-course of sleep/wake states). Each black tick marks a single 4s data epoch. W - wake; N - NREM; R-REM. (C) Representative traces of frontal (F) and parietal (P) EEG (top), EMG (middle), and ACCp GCaMP6s (bottom) signals during REM (left), wake (middle), and NREM (right). For ACCp signal, horizontal gray line represents 0 of the zscored df/f. Black vertical calibration bars in the utmost right represent 1mV (EEG & EMG) and 1std (GCaMP). Black horizontal calibration bar in the bottom right corner represents 1s. (D) Representative scatter plot distribution of EMG root mean square (y-axis) versus frontal EEG power distribution (ratio between power in high [> 25Hz] versus low [< 5Hz] frequencies, x-axis). Each dot marks a single 4s data epoch. Red, wakefulness; Green, REM; Blue, NREM. Wake is associated with high-frequency EEG activity and high muscle tone, NREM is associated with low-frequency EEG activity and low muscle tone, and REM is associated with high-frequency activity and low muscle tone. Embedded pie chart (top left) shows average time spent in each state across the entire data (n=12 mice). (E) Average ACCp claustrum calcium activity in REM, wake, and NREM (n=6). (F) Normalized EEG power (% of total power, y-axis) as a function of frequency (Hz, x-axis) in each state as a function of ACCp claustrum activity (quartiles, n=6). Left, REM (green); Middle, wake (red); Right, NREM (blue). Insets (top right corner) show SWA-to-theta ratios (y-axis) for each ACCp activation quartile (x-axis; from minimal to maximal) in each animal separately (n=6). Mean ratios are depicted as a black line, and individual animals as dashed lines. (G) Representative traces of EEG (top - frontal and parietal), EMG (middle), and ACCp GCaMP (bottom) in auditory stimulation trials associated with maintained sleep (left) vs. awakening (right). Purple vertical bars mark intervals of 1s tone-pip presentation (Methods). Scale bars as in C. (H) ACCp baseline activity (y-axis) for trials associated with maintained sleep (left) vs. awakening (right). Each dot represents a separate ∼10h experiment (11 experiments in n=6 mice). Unless noted otherwise, data are mean ± s.e.m. *p < 0.05, **p < 0.01, ***p < 0.001; n.s., not significant. See Supplementary Table 3 for further details of the statistical analyses.

Given the association between ACCp activity and engagement in the ENGAGE task, together with the tight relation between ACCp activity and SWA (which is associated with the depth of natural sleep), we hypothesized that high ACCp activity levels would also be associated with a deeper disengagement from the sensory environment during sleep, leading to a lower probability of sensory-evoked awakenings. To examine this, we set up an auditory arousal threshold experiment ^7^, where we delivered sounds approximately every minute and determined offline whether each trial resulted in sound-evoked awakening (Figure 5G; Methods). We found that in 9 of 11 recordings, ACCp pre-trial activity was higher before ‘maintained sleep’ trials than before trials resulting in awakening (Figure 5H). Again, this profile was specific for the ACCp network, as OFCp activity did not significantly differ between events leading to awakening vs maintained sleep (Figure S10B). Together, these data establish a specific association between ACCp activity and engagement, where activity in this claustro-cortical projection network is maximal during NREM sleep and in deep sleep when sensory stimuli rarely wake up the animals, and is further associated with higher EEG SWA-theta ratio across behavioral states.

## Discussion

### A unifying perspective on the role of the claustrum

Proposals regarding the function of the claustrum have been framed within the context of two seemingly distinct timescales. On the one hand, the claustrum has been proposed to function in the processing of acute sensory events or distractors, in the context of salience and the gating of sensory perception ^18–21,32,38,46^. On the other hand, claustrum activity has been linked to slow, state-like transitions and oscillations, and even consciousness ^13,14,22,47^. The detailed description provided in this work, of the activity of distinct claustrum populations during the ENGAGE task, as well as during natural sleep, bridges acute and prolonged timescales. Our observations are consistent with the majority of experimental observations and hypotheses published to date regarding the function of the claustrum, and identify a role for ACCp claustral neurons in controlling the full continuum of engagement, providing a holistic and consistent framework for unifying the different perspective on claustral function (Figure S1).

Our study supports the existence of parallel modules of claustrum function, by defining two anatomically, physiologically, and functionally distinct networks, identified by a projection bias to ACC or OFC. Task events and outcomes are broadcast to both OFC and ACC, while trial onset, the most predicable element of the task, was reported selectively by ACCp neurons. Recruitment patterns of projection-defined claustral neurons identified in our recordings may reflect differences in their passive properties ^32,48-50^ or genetic identity ^19,20,36,46,51^, but may also define an orthogonal dimension of behaviorally-relevant ensembles of claustrum neurons.

Whereas OFCp pre-trial activity was uncorrelated with performance, high ACCp pre-trial activity correlated with misses, and low activity correlated with impulsive errors. The significance of these results is enhanced considering previous findings, in which silencing a genetically-defined subpopulation of claustrum projection neurons increased impulsive errors in the presence of sensory load ^19^. It is likely that this effect is mediated primarily by ACCp neurons ^20^. Modulation of the axis of engagement is of broad clinical significance, ranging from hyper-engagement associated with attentiondeficit disorders ^52^, and schizophrenia ^53,54^, to hypo-engagement and apathy, commonly observed in neurodegenerative disorders ^55^. The identification of a specific pathway bi-directionally controlling the full extent of the engagement axis is anticipated to serve as the basis for novel approaches for therapeutic intervention.

The capacity to explore a large number of trials across a rich parameter space within the context of the ENGAGE task, exposed individual differences in attentional strategies employed by mice. Some mice were discriminatory in their approach, such that *selective* mice exhibited a bias towards the visual cues and most prominent auditory cues, while *consistent* mice prioritized the auditory cue over the infrequent visual cue. However, most mice *(erratic)* exhibited behavior that was less moderated, attempting to respond to all the cues within the task (auditory cues of all attenuations, as well as visual cues). The hyper-vigilant approach of *erratic* mice exposed them to impulsive actions in response to the trial onset tone, as well as the tone cloud distractor. This approach also appeared to be exhausting, leading to erratic mice to streaks of missed trials. Intriguingly, the ACCp signal of mice within the *erratic* group exhibited the most prominent representation of the trial onset tone, as well as the strongest bidirectional correlations with impulsive actions and omissions. While it is likely that individual differences in behavioral strategies and corresponding neural signals relate to task engagement, few mechanistic studies have probed these relations^56^. Broader implementation of automated training, supporting the investigation of sophisticated behavioral paradigms in large cohorts of animals, will enable further exploration of individual differences.

Likewise, our results support and shed new light on recent work associating claustrum activity and sleep, by establishing that the high ACCp claustral activity is associated with decreased engagement during NREM sleep, and a reduced probability to awaken in response to auditory stimuli. This is in line with the fact that ablation of the claustrum most prominently impacts SWA in the ACC ^14^. In addition, our long recordings during undisturbed sleep provide the first characterization of claustrum activity during REM sleep, which constitutes less than 10% of the light period in mice ^43^. We found that claustrum activity is predominantly silent during REM sleep, an intriguing observation that is outside the scope of this report, but is worthy of further study.

### Outstanding questions

Evidence so far suggests that widespread activation of claustral projections to ACC would provide a feed-forward inhibitory signal ^19,23^. However the mechanism through which this would modulate engagement remain an open question. The role of the ACCp in gating engagement can be phrased as promoting either response inhibition or attentional inhibition ^57^. On the one hand, the ACC has been implicated in conveying a corollary discharge of the motor plan to sensory areas ^58,59^. Reduced activity of inhibitory signals that regulate this function could lead to premature execution of a motor plan in response to an irrelevant sensory distraction, i.e. the ACCp functions in response inhibition ^60^. On the other hand, the activity of the ACCp network could regulate sensory perception through modulation of sensory cortex, either directly ^19^ or by amplifying cortico-cortical projections from the ACC to sensory cortical targets, i.e. the ACCp functions in attentional selectivity ^23,61^. As perception in rodents is predominantly inferred indirectly via a motor action such as a lick, dissociating between response inhibition and attentional selectivity is challenging. Furthermore, predictive processing and perception may be intimately intertwined ^62^. Resolving the specific mechanistic cognitive function implemented by ACCp neurons remains open for further investigation, likely requiring matched cortical and claustral recordings and manipulations.

Another mechanism through which the claustrum could modulate cortical activity is by impacting cortical synchronicity at different frequencies ^22^. High frequency cortical oscillations have been closely linked to attentional processes and arousal ^45,63,64^. In addition, the claustrum has been implicated in regulating slow waves during sleep, and generating sleep-like dynamics during wakefulness ^13,14,65^. Our results demonstrate a causal role for pre-trial claustrum activity fluctuations in determining performance, as well as a correlation of ACCp activity and SWA-to-theta ratio of cortical EEG. Furthermore, we observe a reduction in ACCp activity during REM sleep, a behavioral state associated with desynchronized cortical activity. It is therefore possible that the mechanism through which the claustrum modulates engagement is by impacting regional cortical oscillations, which differ during vigilant behavior versus lapses ^66^. The limited temporal dynamics of calcium signals do not lend themselves to answer questions relating to fast neural activity and oscillations, and future electrophysiological studies are likely to shed more light on the spectral properties of claustro-cortical activity during behavior.

In summary, ascribing a role for claustral neurons projecting to the anterior cingulate cortex in modulating the full axis of engagement, from hyper-vigilance and impulsivity through to ‘zoning out’ and on to deep sleep, provides a holistic explanation of claustral function. Importantly, this framework implies that the breadth of circumstances during which the claustrum is recruited is wider than previously thought, likely spanning the entire range of events in which salient information is processed, such as learning, rewarding or stressful events, and social behavior.

## Methods

### Animals

All mice described in this study were male C57BL/6JOLAHSD obtained from Harlan Laboratories, Jerusalem Israel. Mice were housed in groups of same-sex littermates and kept in a specific pathogen-free (SPF) animal facility under standard environmental conditions-temperature (20–22°C), humidity (55 ± 10 %), and 12-12 h light/dark cycle (7am on and 7pm off), with ad libitum access to water and food. Mice were randomly assigned to experimental groups. All experimental procedures, handling, surgeries and care of laboratory animals used in this study were approved by the Hebrew University Institutional Animal Care and Use Committee (IACUC; NS-19-15584-3; NS-19-15788-3). While all experiments were performed in male mice, we do not anticipate the results to differ drastically between males and females.

### Surgery

#### Stereotactic surgery and viral injections

Induction and maintenance of anesthesia during surgery was achieved using SomnoSuite Low-Flow Anesthesia System (Kent Scientific Corporation). Following induction of anesthesia, animals were quickly secured to the stereotaxic apparatus (David KOPF instruments). Anesthesia depth was validated by toe-pinching and isoflurane level were adjusted (1-5%) to maintain a heart rate of ∼60bpm. The skin was cleaned with Betadine (Dr. Fischer Medical), Lidocaine (Rafa Laboratories) was applied to minimize pain, and Viscotear gel was applied to protect the eyes. An incision was made to expose the skull, which was immediately cleaned with Hydrogen peroxide (GADOT), and a small hole was drilled using a fine drill burr (model 78001RWD Life Science). Using a microsyringe (33GA; Hamilton syringe, Reno, NV) connected to an UltraMicroPump (World Precision Instruments, Saratosa, FL) virus was subsequently injected at a flow rate of 50-100nl/min, following which the microsyringe was left in the tissue for 5-10 minutes after the termination of the injection before being slowly retracted. For photometry experiments, an optic fiber ferrule (400um, 0.37-0.48 NA, Doric Lenses) was slowly lowered into the brain. A custom-made metal head bar was glued to the skull, the incision was closed using Vetbond bioadhesive (3M) and the skull was covered in dental cement and let dry. An RFID chip (ID-20LA, ID Innovations) was implanted subcutaneously. Mice were then disconnected from the anesthesia, and were administered with subcutaneous saline injection for hydration and an IP injection of the analgesic Rimadyl (Norbrook) as they recovered under gentle heating. Coordinates for the claustrum were based on the Paxinos and Franklin mouse brain atlas ^67^. Unless noted otherwise, viruses were prepared at the vector core facility of the Edmond and Lily Safra Center for Brain Sciences, as described previously ^26^.

**Supplemental Table T1:**
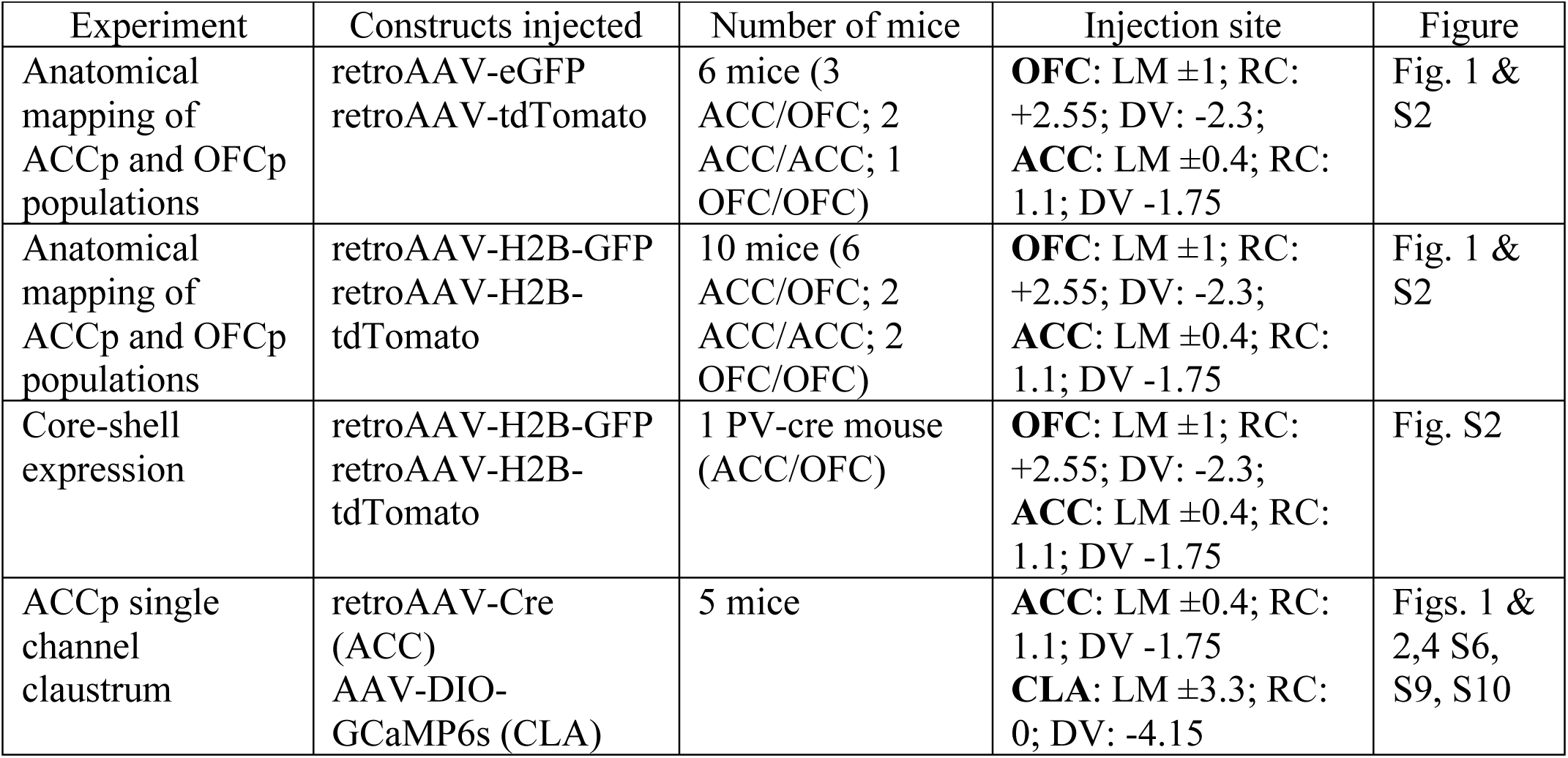
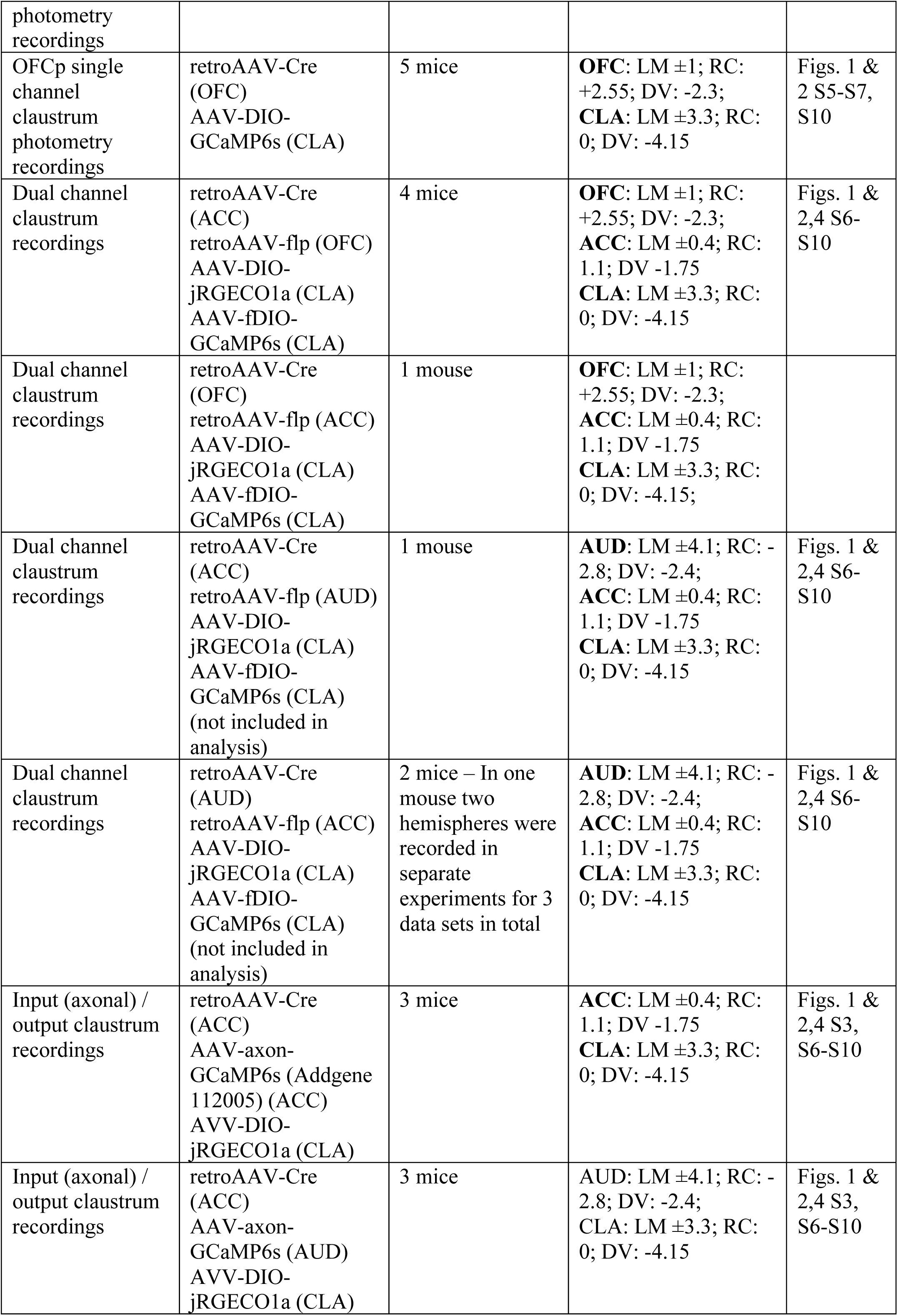
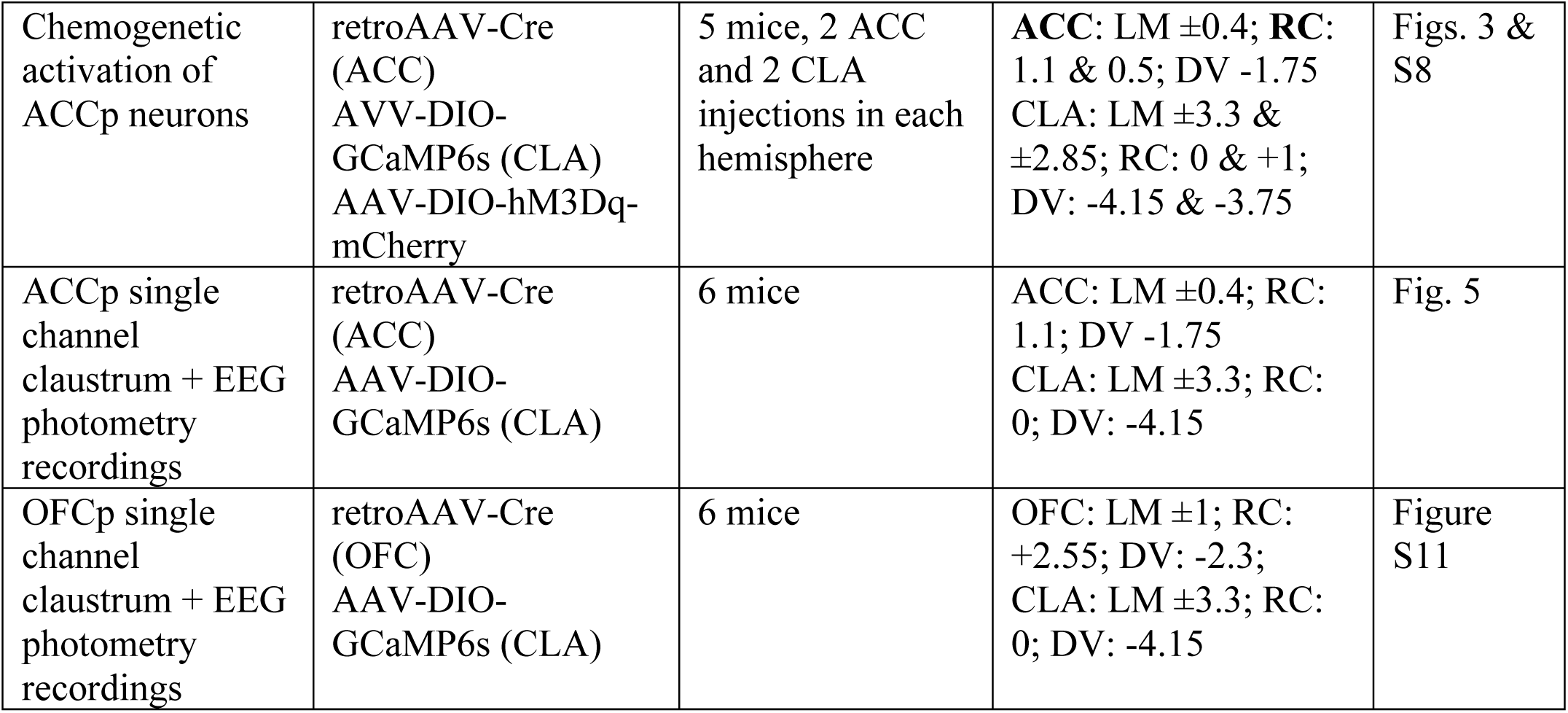
Surgeries

#### EEG and EMG

12 mice underwent stereotactic surgery for viral expression of GCaMP in claustral projection neurons [retro-AAV-CRE from ACC and DIO-GCaMP6s in the claustrum (n=6); retro-AAV-CRE from OFC and DIO-GCaMP6s in the claustrum (n=6)], implanted with a fiber over the left claustrum and prepared for EEG and EMG recordings. Two screws, frontal and parietal (1 mm in diameter) were placed over the right hemisphere for EEG recording. Two additional screws were placed above the cerebellum as reference and ground. Two single-stranded stainless-steel wires were inserted to either side of the neck muscles to measure EMG. EEG and EMG wires were soldered onto a custom-made headstage connector. Dental cement was used to cover all screws and EEG/EMG wires. Following validation of photometry signal, mice were transported to Tel Aviv University for further recordings.

#### Histology

Mice were anesthetized for terminal perfusion by a mix of Ketamine/Xylazine and perfused with cold PBS, followed by 4% PFA. Following decapitation, heads were placed in 4% PFA overnight to preserve the location of the optic ferrule. Brains were then carefully extracted, and placed in 4% PFA for another night prior to transiting to PBS in preparation for sectioning and histology. The fixed tissue was sectioned using a Vibratome (7000 smz-2) at 60μm thickness.

In order to enhance GCaMP6s signals for analysis of axonal projections, floating section immunohistochemistry was performed (rabbit anti-GFP, Life Technologies, Bethesda, MD; catalog No. A-6455; final dilution to 1:500 in 3% normal horse serum), following previously described protocols ^26^. mCherry (jRGECO) signal was likewise amplified for the visualization of DREADD expression (rabbit anti-RFP; Rockland, Limerick, PA; catalog No. 600-401-379; final dilution 1:1000 in 3% normal horse serum).

#### Image acquisition

Slides were scanned on a high-speed fully-motorized multi-channel light microscope (Olympus IX-81) in the microscopy unit of the Alexander Silberman Institute of Life Sciences. Slices were imaged at 10X magnification (NA=0.3), green and red channels exposure times were selected for optimal clarity and were kept constant within each brain series. DAPI was acquired using excitation filters of 350±50 nm, emission 455±50 nm; eGFP excitation 490±20 nm, emission 525±36 nm; tdTomato and Ruby excitation 555±25 nm, emission 605±52 nm and for Alexa 647 excitation 625 nm, emission 670 nm.

#### Quantification and statistical analysis

Cell counting and co-localization analysis: In order to quantify labelled cells (number and overlap), automated image analysis was used. For each claustrum seven images were captured from Bregma 1.1 mm to Bregma -1.06 mm, from 60 μm thick slices separated by 360 μm. The claustrum was manually cropped according to the outline depicted in the appropriate section from the Paxinos and Franklin mouse brain atlas ^67^. The image files of the cropped claustrum were used in the analysis pipeline, three channels for each image: DAPI, eGFP and tdTomato or Ruby. The data was analyzed using the CellProfiler v.3.0.0 co-localization pipeline (www.cellprofiler.org), with minor modifications, including feature enhancement and shrink/expand objects ^68,69^. For fluorescently labelled retroAAV analysis (n = 3 ACC/OFC; 2 ACC/ACC OFC/OFC mice), a DAPI object mask was generated and objects from eGFP and tdTomato channels that overlap with the mask were considered labelled cell bodies. Overlap was defined as the overlay of a detected cell body from the GFP channel coinciding with a cell body in the tdTomato channel, both coinciding with the DAPI mask. RetroAAV-H2B (6 ACC/OFC; 2 ACC/ACC; 2 OFC/OFC mice) expressed in the nuclei, providing lower background and allowing detection of labelled nuclei directly from eGFP and Ruby channels without a DAPI object mask. In addition, the analysis was modified such that object centroid distances were measured and calibrated such that only objects with a maximal 6 pixel centroid distance between them were considered to be double-labelled cells. Histograms corresponding to the spatial localization of the labelled cells were built in RStudio (Ver. 1.0.153).

#### Quantification of projections

Axonal projections of ACCp and OFCp populations were quantified from sections obtained from brains of mice which participated in the task (ACCp - 4 mice; OFCp - 3 mice), immuno-stained to enhance indicator (GCaMP6s or jRGECO1a, see above) fluorescence in projections. After alignment of section images to the Paxinos and Franklin mouse brain atlas ^67^ a manual threshold was set for every brain such that the claustrum area would be saturated, and background minimal, enabling a clear contrast for fluorescent processes. Structures of interest were selected based on previous anterograde tracing studies in the claustrum ^27^. Analysis was conducted in Fiji (ImageJ) and quantified as the mean pixel intensity in a rectangle sampled within different brain divisions. Measurements were obtained from qualitatively similar positions in each section across mice.

#### Automated behavioral training

Training cages comprised of a 4cm diameter tube corridor connected to the home cage of the mice. At the end of the corridor a behavioral lick port (Sanworks) was positioned. Within the training cage mice had ad libitum access to food, while access to water was restricted to the output of the behavioral system. A radio-frequency identification (RFID) reader (ID-20LA, ID Innovations) was positioned above the corridor for individualized identification of mice. Auditory cues were delivered by a Bpod wave player (Sanworks) connected to earphones positioned on the corridor adjacent to the port. Experiments were controlled via an open source code MATLAB-based state machine (Bpod, Sanworks). A custom protocol was written in MATLAB in order to support individualized training by gating the Bpod state machine as a function of the output of the RFID reader. This enabled activation of different task parameters for individual mice based on their performance. Training comprised several stages, and each mouse progressed individually, according to its learning. Mice were then taught to associate the auditory-visual cue with water availability during a lick adaptation period. Entry of a mouse into the port (an RFID reading) initiated a trial, reported to the mouse by a 0.1 sec broadband noise (BBN, intensity = 70.5db SPL) marking trial onset. Trial initiation was followed by a varying delay period in which mice had to withhold lick responses. This delay period lasted 0.1 sec in the adaptation phase and was prolonged in the following training steps. If the mouse successfully withheld licking, a cue was presented at the end of the delay period, consisting of 5 pure tone pips of 6 kHz, 0.1 sec long (spaced 0.1 sec, intensity = 86.1db), a white LED light (these auditory/visual cues were referred to as AudVis). The first lick within a 1.5s window following cue onset was rewarded (10ul of drinking water). Impulsive or late licks were not rewarded, and mice had to exit the port (terminate and reinitiate RFID reading) before a new trial could be initiated. After mice reached satisfactory success rates (50-70% correct, 2.6 days on average) they proceeded to stage 2 where the delay was prolonged to between 0.5-2s (2.5 days on average). Mice proceeded to stage 3, which included the full range of possible delays (0.5-3s) and a gradual transition to auditory trials with no visual aid (Aud) in three steps: 30% Aud (Stage 3a), 50% Aud (Stage 3b), and 70% Aud (Stage 3c). Following stage 3 (4.4 days on average) a pure tone-cloud masking stimulus was introduced (4s of continuous chords assembled from logarithmically spaced pure tones in the frequency range of 1-10kHz, excluding the target cue frequency, intensity = 67.5db SPL), lasting from trial onset throughout the delay and cue. The tone-cloud was also introduced gradually. Stage 4a comprised of 70% auditory-visual trials with tone cloud (AudVisCloud) and 30% Aud (2.7 days in average). Stage 4b included 50% AudVisCloud trials, 20% auditory cloud trials (AudCloud), 15% Aud trials and 15% AudVis trials. After mice were familiar with the cloud in both visual and non-visual trials, we proceeded to stage 4c, and increased the rate of cloud trials to 65% AudCloud, while the rest of the trials comprised 15% AudVisCloud, 15% Aud, and 5% AudVis (mice spent on average 4.5 days in stages 4b + 4c). Finally, we gradually added 3 attenuations of the target cue. First, in stage 5a, 30% of the trials included the full range of attenuations (Go-Cue trial intensities were (db SPL): #1: 68.75db; #2: 81.2db; #3: 86.1db; #4: 91.6db), which increased in stage 5b to 50% of the trials and then in stage 5c to 100% percent of the trials (13 days on average, depending on the availability of the recording system, adding up to a mean total of 29.6 days of training). Response duration and reward size were kept constant throughout training. Due to the COVID-19 pandemic, the training schedule of two mice was altered, and they were thus excluded from panels illustrating training data.

Task structure during head-restrained recordings was identical to the automated training, except that trials were initiated automatically every 20 seconds. Behavioral sessions contained blocks of trials containing 15 (in some cases shortened to 8 or 10) occurrences of each possible combination of parameters, in random order. These blocks were repeated 2-4 times as long as mice maintained participation, for a total of up to 1000 trials / mouse / day (sessions typically extended over 240-480 trials). Trials in which the mouse licked late (>1.5s) were rare (∼2% of all trials) and were thus also labelled as misses. The degree to which different trial parameters (cue intensity, cloud, visual aid) affected behavior was quantified by calculating a modulation index for the effect of each parameter on hit rates in the task (i.e. difference normalized by sum: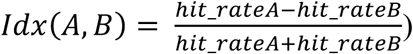. For cue modulation, we compared hit rates in the second lowest intensity, which was the most variable, to the strongest intensity.

#### *In Vivo* fiber photometry recordings

Fiber photometry data was collected using a 1-site Fiber Photometry system (Doric Lenses, Canada) adapted to two excitation LEDs at 465nm (calciumdependent GCaMP fluorescence) and either 405nm (isosbestic control channel) or 560nm (for two-color recording using jRGECO). Simultaneous monitoring of the two channels was made possible by connecting the LEDs to a minicube (with dichroic mirrors and cleanup filters to match the excitation and emission spectra; FMC4 or FMC5, Doric) via an attenuating patch cord (400 μm core, NA=0.37-0.48). LEDs were controlled by drivers that sinusoidally modulated 560nm/465nm/405nm excitation at 210/210/330Hz, respectively enabling lock-in demodulation of the signal (Doric Lenses, Canada). Zirconia sleeves were used to attach the fiber-optic patch cord to the fiber implant on the animal. Data were collected using Femtowatt photoreceiver 2151 (Newport) and demodulated and processed using an RZ2 (at TAU) or RZ5P (at HUJI) BioAmp Processor unit and Synapse software (TDT). LED intensities were individually modulated in each mouse to allow the recording of viable signals with the minimal intensity possible. To this end, 465nm LED intensity was gradually increased until robust GCaMP/jRGECO fluctuations were observed above noise. Next, the 405nm (isosbestic control channel) LED intensity was set to allow detection of motion artifacts. The total power at the tip of the patch cable was most often 0.05-0.1mW. The signal, originally sampled at 24414Hz, was demodulated online by the lock-in amplifier implemented in the processor, sampled at 1017.25Hz and low-pass filtered with a corner frequency at 4Hz. All signals were collected using Synapse software (TDT). EEG and EMG were digitally sampled at 1017 Hz (PZ2 amplifier, Tucker-Davis Technologies), and filtered online: both signals were notch filtered at 50/100 Hz to remove line noise and harmonics; then, EEG and EMG were band-pass filtered at 0.5-200Hz, and 10-100Hz, respectively. Due to a technical issue, EEG were also high-pass filtered in hardware > 2Hz but a comparison with full broadband (>0.5Hz) EEG in several animals verified signal differences were minor and did not affect the ability to analyze sleep stages or SWA. Simultaneous video data (used for sleep scoring and for behavioral assessments) were captured by a USB webcam (at TAU) or an IR camera (at HUJI, Basler) synchronized with electrophysiology/photometry data. Offline, EEG and EMG were resampled to 1000 Hz (MATLAB, The MathWorks) for sleep scoring and power spectrum analysis.

Behavioral fiber photometry recordings were made in one of three head-restrained conditions: 1) Spontaneous recordings, in which no stimuli were presented, and the mouse was free to run on a treadmill (Janelia 2017-049 Low-Friction Rodent-Driven Belt Treadmill) for 10min (for validation of chemogenetic effects) or 40min (for correlation analyses). 2) Passive auditory sessions, in which broadband noise or frequency sweeps (1-40Khz played at a 100kHz sample rate, through an RP2.1 processor, TDT), attenuated between 0-20db (SA1 amplifier, PA5 attenuator, TDT) were played while the mouse was free to run on a treadmill. 3) Task sessions, which consisted of several blocks (1-4), each consisting of 120-180 trials, as described above.

#### Fiber photometry analysis

Unless otherwise noted, all analysis was performed using custom MATLAB scripts. First, to correct for baseline drift due to slow photobleaching artifacts, particularly during the first several minutes of each session, a 5^th^ order polynomial was fit to the raw data and then subtracted from it. After baseline correction, ΔF/F was computed using the 99^th^ lowest percentile value as 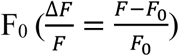, and the resulting trace was z-scored relative to the mean and standard deviation of the entire recording session to normalize between channels and across mice. For 2/30 mice, motion artifacts were corrected by using the z-scored isosbestic control channel as a sample-by-sample F^0^ for computing AF.

To correct for small session-to-session fluctuations in the signal, while maintaining quantitation of pretrial activity, we calculated pre-trial activity for every individual trial (four seconds before trial onset), and used the pre-trial signal as a dependent variable in a linear model with recording session and trial outcome as independent variables (baseline ∼ outcome + session). A scalar value of the intercept and estimate for each session was then subtracted from the corresponding data set, setting the mean baseline for correct trials for each session at approximately zero. Pre-processed data was then cut into 20 second windows (−5:15 seconds) around each behavioral epoch: trial onset, cue onset, lick onset and run onset, and concatenated for each mouse to form an event-aligned activity matrix together with an information table detailing the parameters and outcome of each trial.

#### Analysis of spontaneous claustrum activity

Spontaneous calcium events were identified with the MATLAB function *findpeaks*. To avoid multiple identification of single events or defining noise as activity, we employed a threshold of a minimum prominence of 1 standard deviation and a minimum of 2 seconds event width, measured at half prominence. Changing these parameters did not drastically alter results.

#### Linear encoding model

A linear encoding model was constructed, using ridge-regression to create time-averaged kernels for each behavioral epoch in the task, following code from Musall et al., 2019 ^41^.For each mouse, all trials from all sessions were used to create the full model. Ten-fold cross validated estimation of the explained variance by the full model (CVR^2^) was then compared to that explained by each individual label on its own *(single variable model)*. The *unique contribution* of each variable was estimated by the loss in explained variance (AR^2^) by omitting each variable from the full model. Both measures were normalized to the size of the window (Supplemental Table T2) and the CVR^2^ of the full model. Estimation statistics (www.estimationstats.com) were performed based on the work described in ^70^.

**Supplemental Table T2:**
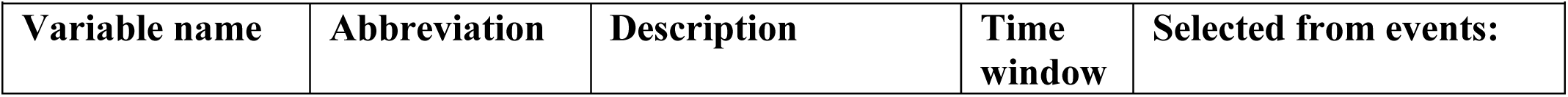
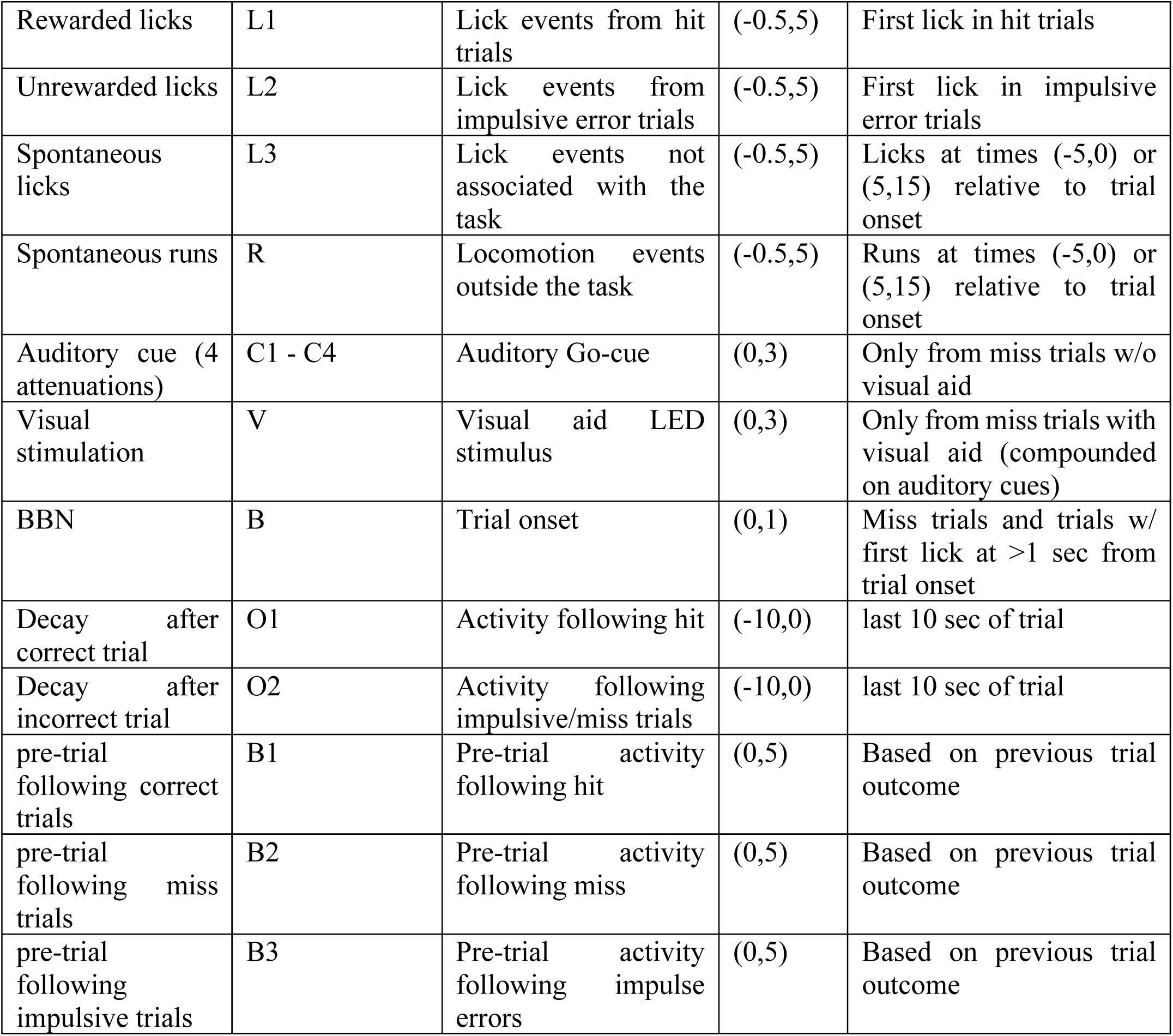
Epochs for time-event kernels in the linear encoding model.

|  |  |  |  |  |
| --- | --- | --- | --- | --- |
| Variable name | Abbreviation | Description | Time window | Selected from events: |
| Rewarded licks | L1 | Lick events from hit trials | (-0.5,5) | First lick in hit trials |
| Unrewarded licks | L2 | Lick events from impulsive error trials | (-0.5,5) | First lick in impulsive error trials |
| Spontaneous licks | L3 | Lick events not associated with the task | (-0.5,5) | Licks at times (-5,0) or (5,15) relative to trial onset |
| Spontaneous runs | R | Locomotion events outside the task | (-0.5,5) | Runs at times (-5,0) or (5,15) relative to trial onset |
| Auditory cue (4 attenuations) | C1 - C4 | Auditory Go-cue | (0,3) | Only from miss trials w/o visual aid |
| Visual stimulation | V | Visual aid LED stimulus | (0,3) | Only from miss trials with visual aid (compounded on auditory cues) |
| BBN | B | Trial onset | (0,1) | Miss trials and trials w/ first lick at >1 sec from trial onset |
| Decay after correct trial | O1 | Activity following hit | (-10,0) | last 10 sec of trial |
| Decay after incorrect trial | O2 | Activity following impulsive/miss trials | (-10,0) | last 10 sec of trial |
| pre-trial following correct trials | B1 | Pre-trial activity following hit | (0,5) | Based on previous trial outcome |
| pre-trial following miss trials | B2 | Pre-trial activity following miss | (0,5) | Based on previous trial outcome |
| pre-trial following impulsive trials | B3 | Pre-trial activity following impulse errors | (0,5) | Based on previous trial outcome |

#### Chemogenetic activation

30 minutes prior to each recording session, mice received an IP injection of either saline as a control or clozapine-n-oxide (CNO), diluted to a final dilution of 1mg/ml (10 mg CNO in 500ul DMSO and 9.5 ml saline) and administered at a dose of 10 mg/kg.

#### Spectral analysis of pre-trial dynamics

Pre-trial activity was analyzed over individual sessions of 300 trials each (shorter sessions were excluded from the analysis, and longer sessions were analyzed only up to trial 300). A fast Fourier Transform (using the *fft* function in MATLAB) was applied to each session, and the average power spectrum over sessions was compared to a threshold defined by the maximal power obtained in each frequency over 1000 shuffling iterations of the data. The reported fluctuation frequency is the peak of the power spectrum that crosses this threshold.

#### EEG recordings during natural sleep

##### Undisturbed sleep

Several weeks after surgery (due to transport to TAU), mice (n=12) were placed in a new home cage within an acoustic chamber (40dB attenuation, H.N.A, Israel) and connected to the EEG/EMG headstage and to the optic fiber patch cord through a rotary joint commutator. After > 72h of habituation to the new cage and to tethered recording, electrophysiology and photometry data were recorded continuously for 12 hours during light-phase daytime hours while animals were undisturbed and behaving freely. To minimize bleaching and phototoxicity, LEDs were automatically disengaged for 30 minutes every 2 hours (90min ON/30min OFF).

##### Auditory arousal threshold experiments

Experiments (lasting on average ∼10 hours, starting shortly after light onset) were conducted in a double-wall sound-attenuating acoustic chamber (Industrial Acoustics Company, Winchester, UK). Sounds were generated in TDT software, amplified (SA1, Tucker Davis Technologies (TDT), and played free-field through a magnetic speaker (MF1, TDT) mounted 50cm above the animal. Sound intensities were measured by placing a Velleman DVM805 Mini Sound Level Meter at the center of the cage floor. In arousal threshold experiments, broadband noise bursts (1s duration, either 65dB or 80dB SPL, order counterbalanced) were presented intermittently every 60s (± 0.5 jitter) when mice (n=12) were undisturbed. The sensitivity of the setup was confirmed by verifying that awakening probability was significantly higher for louder sounds (19.8 ± 8.4% vs. 8.1 ± 2.6% for 80dB vs. 65dB SPL sounds, respectively, p<0.001, paired t-test). The analysis presented in Figure 5H is based on the louder sound, for which there was a sufficient number of trials in both conditions (maintained sleep and awakening). Whenever COVID19 lockdown restrictions allowed (n=9/12 animals), we performed two separate experimental sessions per animal.

##### Data and code availability

Full data and code used for creating the figures will be uploaded to a public repository prior to publication.

## Acknowledgments

The authors would like to thank Dr. Rylan Larsen (Allen Institute), for generously gifting us an initial sample of AVV-fDIO-GCaMP6s, and Ege Yalcinbas (UCSD) for directing us towards the axon-targeted GCaMP. We also wish to thank Dr. Simon Musall (UCSD) for providing accessible code, allowing us to implement the linear encoding model. The authors appreciate the helpful critical comments of colleagues, members of the Citri lab, Gal Vishne, Dr. Eran Lottem and Profs. Inbal Goshen, Leon Deouell, Ayelet Landau, Adi Mizrahi and Mickey London on data, writing and presentation. Work in the Citri laboratory is funded by the European Research Council (ERC 770951), The Israel Science Foundation (1062/18, 393/12, 1796/12, and 2341/15), The Israel Anti-Drug Administration, EU Marie Curie (PCIG13-GA-2013-618201), the National Institute for Psychobiology in Israel, Hebrew University of Jerusalem Israel founded by the Charles E. Smith family (109-15-16), an Adelis Award for Advances in Neuroscience, the Brain and Behavior Foundation (NARSAD 18795), German-Israel Foundation (2299-2291.1/2011), and Binational Israel-United States Foundation (2011266), the Milton Rosenbaum Endowment Fund for Research in Psychiatry, a Prusiner-Abramsky Research Award in Basic Neuroscience, a seed grant from the Eric Roland Fund for interdisciplinary research administered by the ELSC, contributions from anonymous philanthropists in Los Angeles and Mexico City, as well as research support from the Safra Center for Brain Sciences (ELSC) and the Canadian Institute for Advanced Research (CIFAR). The Nir lab is supported by the European Research Council (ERC-2019-CoG 864353), the Israel Science Foundation (ISF, grants 1326/15 & 51/11 I-CORE cognitive sciences), and the Adelis Foundation.

## Figure Legends

**Figure S1.**
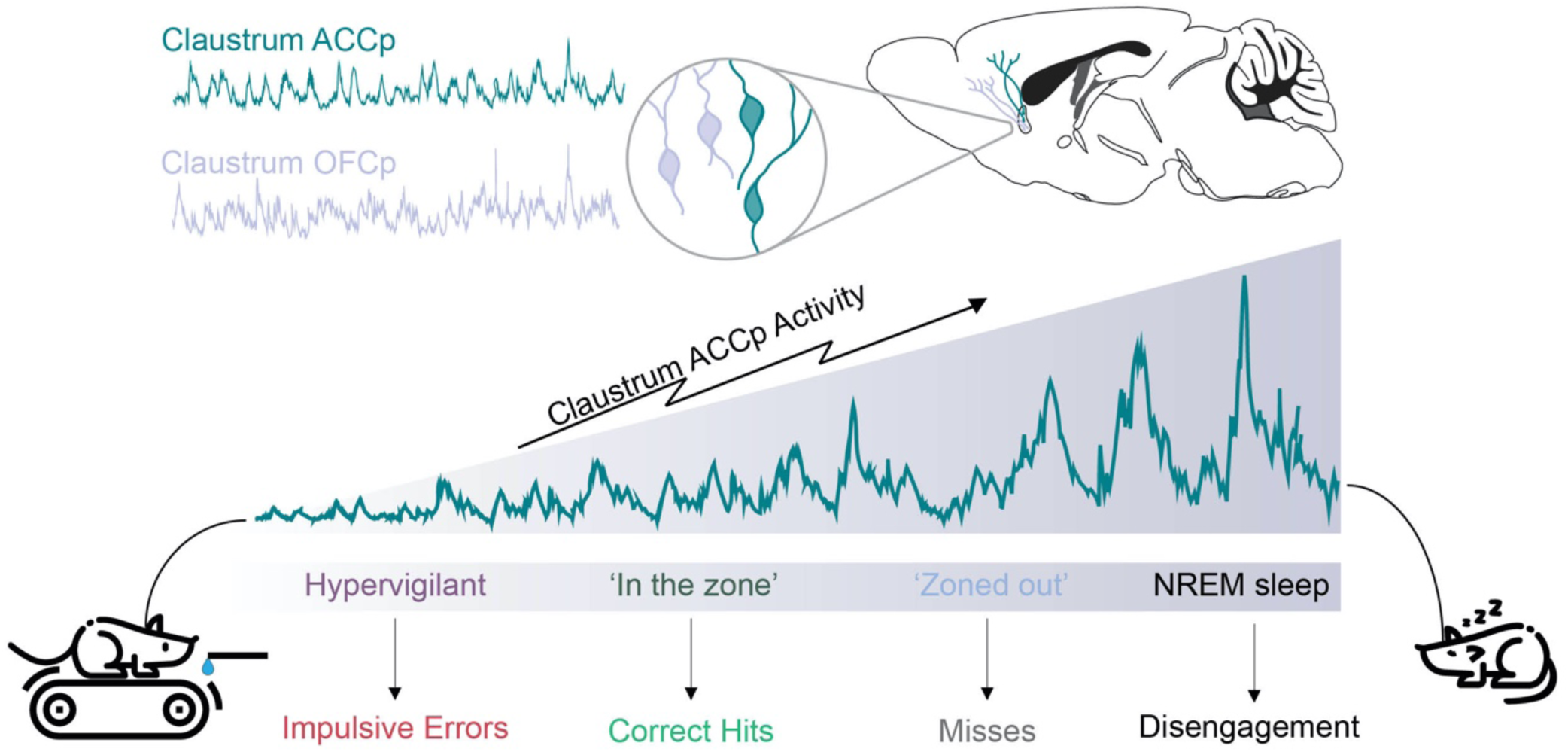
Increased ACCp activity is associated with reduced engagement during behavior and in sleep. Optimal ‘in the zone’ performance requires a defined, moderate, level of ACCp activity (Figures 2-4). At low ACCp activity levels, mice tend to perform impulse errors in the response to the trial onset BBN, rather than withhold their response in anticipation of the ‘go’ cue. At high ACCp activity levels, mice tend to ‘zone out’ and miss trials. Furthermore, even higher levels of ACCp activity are associated with ‘miss streaks’, in which the mice do not engage with the task over multiple minutes. Finally, during sleep, cortical slow-wave EEG is correlated with increased ACCp activity, and the propensity of mice to awake from NREM sleep following tone stimulations decreases as a function of ACCp activity (Figure 5).

**Figure S2.**
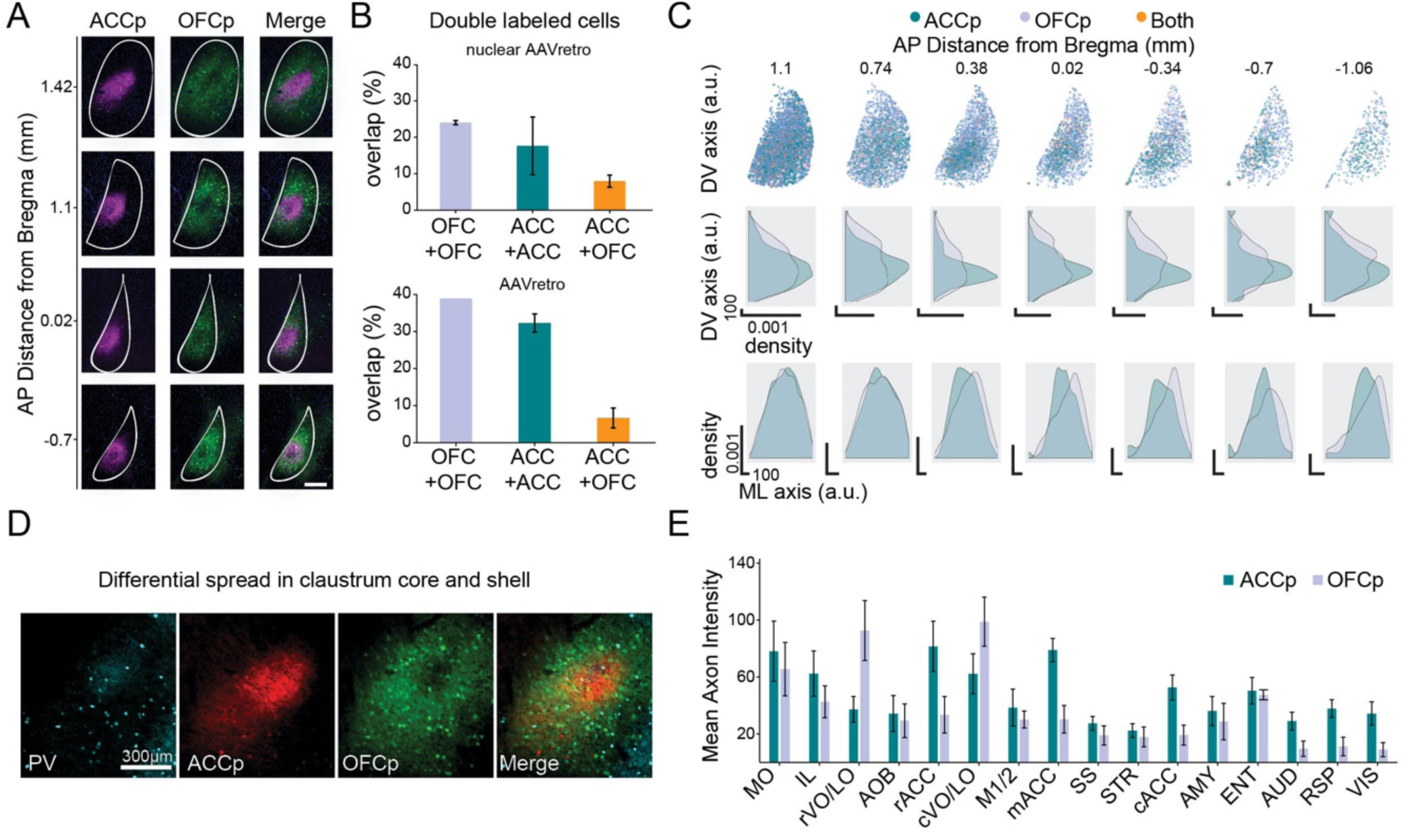
Anatomical distinction of ACCp and OFCp claustral populations. (**A**) Differential localization of claustral ACCp and OFCp networks (labeled by retro-labeling ACC-vs OFC-projecting cells with cytoplasmic tdTomato vs eGFP expression) in several planes along the anterior-posterior claustrum axis. Scale bar represents 100μm. (**B**) Quantification of overlap (percent of all labelled cells) using nuclear-localized (H2B-fused; top) or cytoplasmically-diffuse (bottom) expression of fluorophores driven by retro-AAVs co-injected to the same cortical target vs. different cortical regions. (**C**) Data overlaid from 7 mice showing spread of ACCp and OFCp neurons (top panels) and their respective density distributions along the dorso-ventral (DV; middle panels) or medio-lateral (ML; lower panels) axes of the claustrum. (**D**) Localization of ACCp neurons in the claustrum core, identified by parvalbumin (PV) immunostaining, vs relatively sparse distribution of OFCp network signal within this ‘core’ patch. (**E**) Quantification of axonal intensity by brain region for ACCp or OFCp neurons following anti-GFP immunostaining (expanded data from Figure 1E, see methods). Glossary: MO - Medial orbital cortex; IL - Infralimbic cortex; rVO\LO - ventral orbital cortex, rostral; AO - Anterior olfactory nucleus; rACC - Anterior cingulate cortex, rostral, cVO/LO - Orbitofrontal cortex, caudal; M1/2 - Motor cortex; mACC - Anterior cingulate cortex, middle; SS - Somatosensory cortex; STR - Striatum; cACC - Anterior cingulate cortex, caudal; AMY - amygdala; ENT - entorhinal cortex; AUD - Auditory cortex; RSP - Retrosplenial cortex; VIS - Visual cortex. Data in Figure 1E shows averaged axonal density in ACC (r, m, and c), OFC (r and c VO\LO), and sensory cortex (AUD, RSP and VIS).

**Figure S3.**
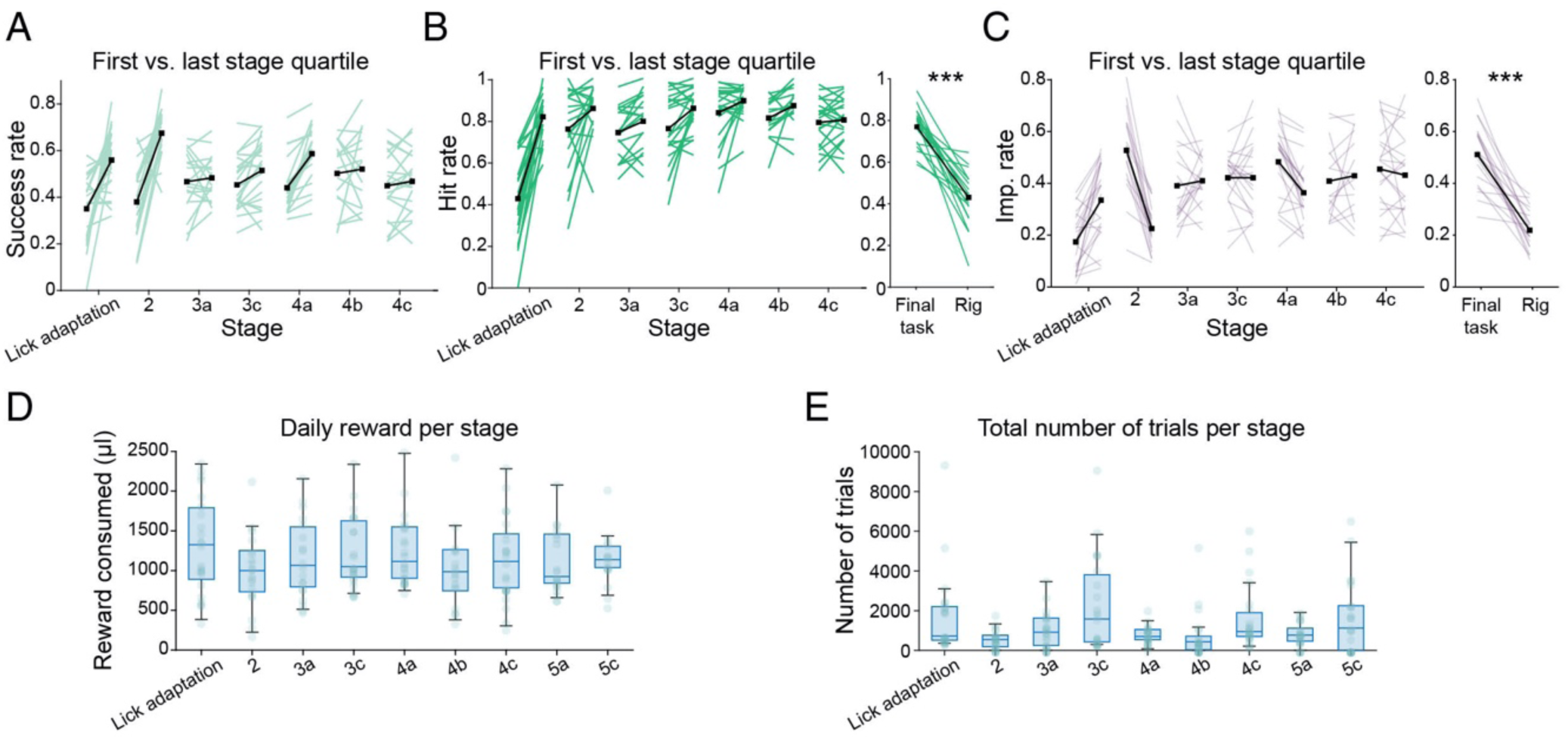
Performance metrics during automated behavioral training. (**A**) Overall success rate during automated training. Points represent performance in the first and last quartiles of each stage. Black lines represent group averages. Training stages proceed from lick adaptation (cue initiated upon port entry; 1.5 second reward window; all trials include visual aid - AudVis); Stage 2: addition of delay prior to cue (random delay of 0.5-2 sec); Stage 3: lengthened delay (0.75-3s) and gradual increase in difficulty by removal of visual aid in three steps, 30% (Stage 3a), 50% (Stage 3b), and 70% (stage 3c) trials are purely auditory (Aud). In stage 4 a tone-cloud distractor was added. Stage 4a comprised of 70% auditory-visual trials with the cloud (AudVisCloud). Stage 4b included 50% AudVisCloud trials, 20% auditory cloud trials (AudCloud), 15% Aud trials and 15% AudVis trials. After mice were familiar with the cloud in both visual and non-visual trials, we proceeded to stage 4c, and increased the rate of AudCloud trials to 65%, while the rest of the trials comprised of 15% AudVisCloud, 15% Aud, and 5% AudVis. Finally, in stage 5, 3 additional attenuations of the target cue were introduced. In stage 5a to 30% of the trials, in stage 5b to 50% of the trials, and in stage 5c to 100% percent of the trials (success rate in the full task during training is shown in Figure 2C). (**B**) As in **A**, for hit rate (excluding impulsive trials). Right panel summarizes the hit rate in the full task during training compared to the head-fixed recordings. The increase in missed trials reflects the change from a self-paced task to a constant 20 second inter-trial interval. (**C**) As in **A, B** for impulsive errors, which were far less prominent during head-fixed recordings (potentially reflecting reduced competition for the port compared to the group-housed automated training). (**D**) Daily reward consumption during training stages. (**E**) Number of trials in each stage. Box plots in (**D**) and (**E**) represent group median and 1st and 3rd quartiles. Dots represent individual mice. Unless noted otherwise, data are mean ± s.e.m. *p < 0.05, **p<0.01, ***p< 0.001; n.s., not significant. See Supplementary Table 3 for further details of the statistical analyses.

**Figure S4.**
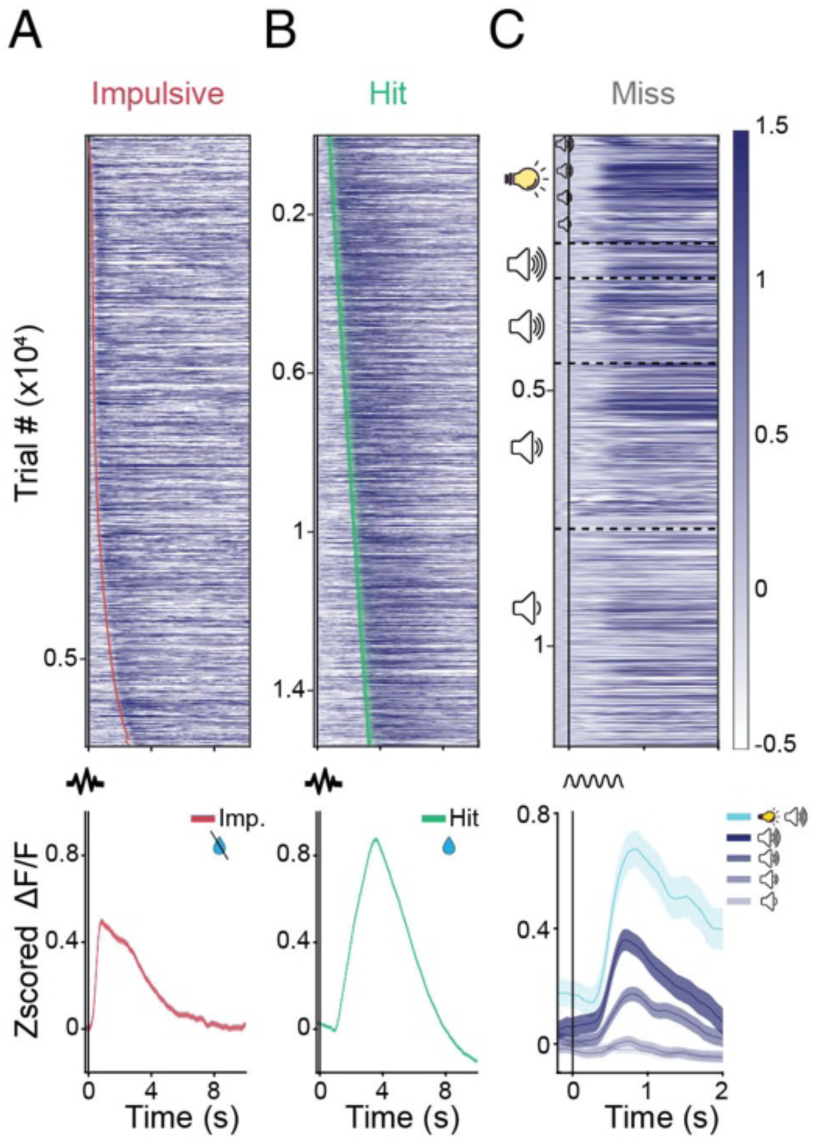
OFCp activity is recruited following licks and cues but not trial onset. Top: All OFCp trials for impulsive (**A**, n=5,835), hit (**B**, n=15,409), and miss (**C**, n=12,815) trials from n=10 mice, sorted according to lick onset (impulsive); delay from trial onset to cue (hits); or cue intensity (miss). Red and green ticks indicate the first impulsive or correct lick within the trial, respectively. Bottom: mean activity traces in impulsive (left) hit (middle) and miss trials (right, separated by cue intensity). Shaded area represents SEM. The vertical black line indicates trial onset (for impulsive & hit trials) or cue (for miss trials).

**Figure S5.**
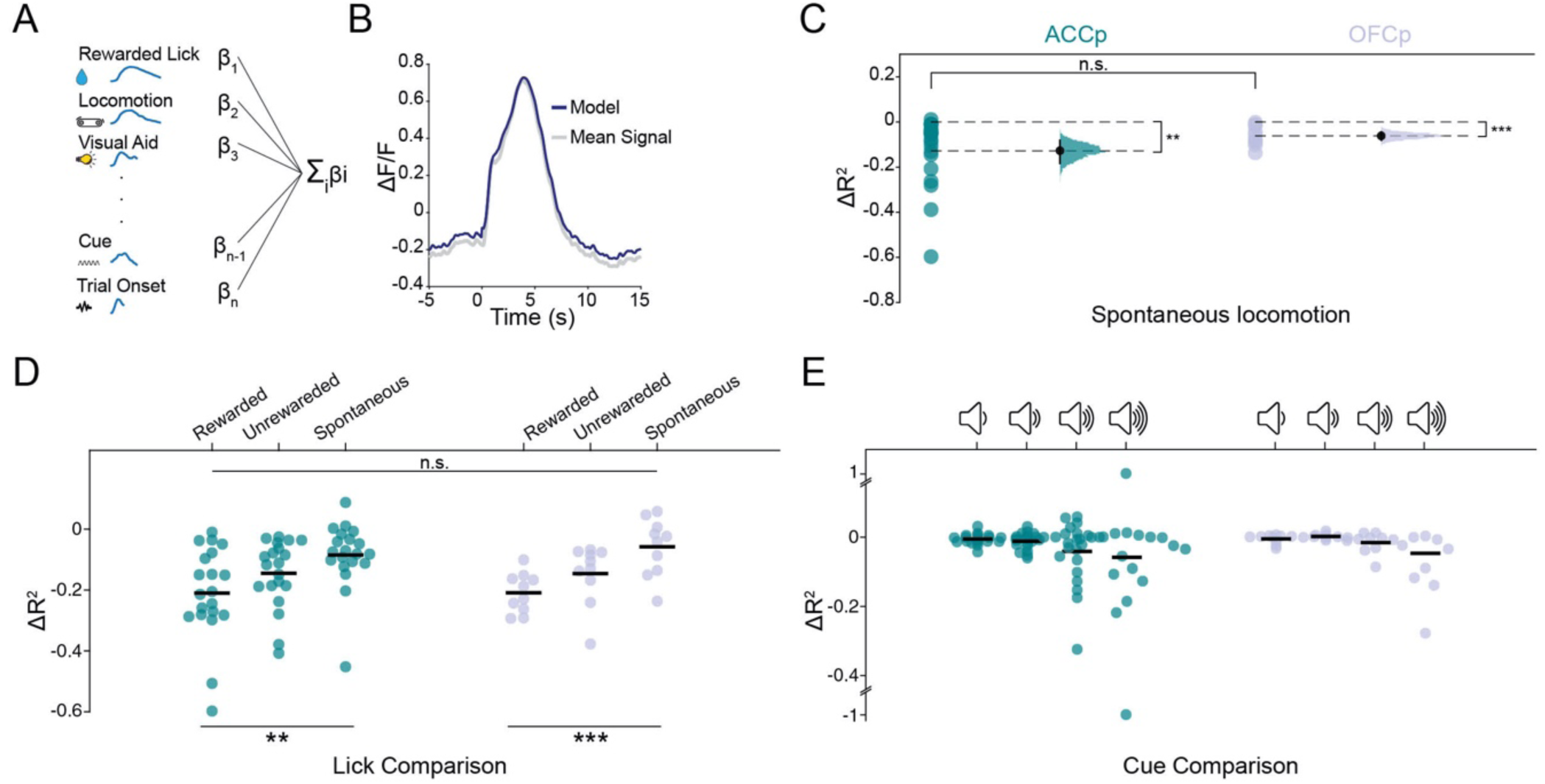
A linear encoding model for quantification of the claustral representation of task parameters. Time-event kernels (**A**) are linearly summed to generate a prediction for the average neural signal (**B**). See Figure 2 and supplementary table T2 for the full list of labels. (**C**) Model quantification of the unique contribution of claustrum activity during spontaneous locomotion events (ACCp n=20; OFCp n=10). Data is shown as individual channels and bootstrapped distribution of means with 95% confidence intervals. (**D-E**) Model quantification of the unique contribution of claustrum activity during licking events (**D**) and go-cue stimuli (**E**) in ACCp and OFCp signals. Unless noted otherwise, data are mean ± s.e.m. *p < 0.05, **p < 0.01, ***p < 0.001; n.s., not significant. See Supplementary Table 3 for further details of the statistical analyses.

**Figure S6.**
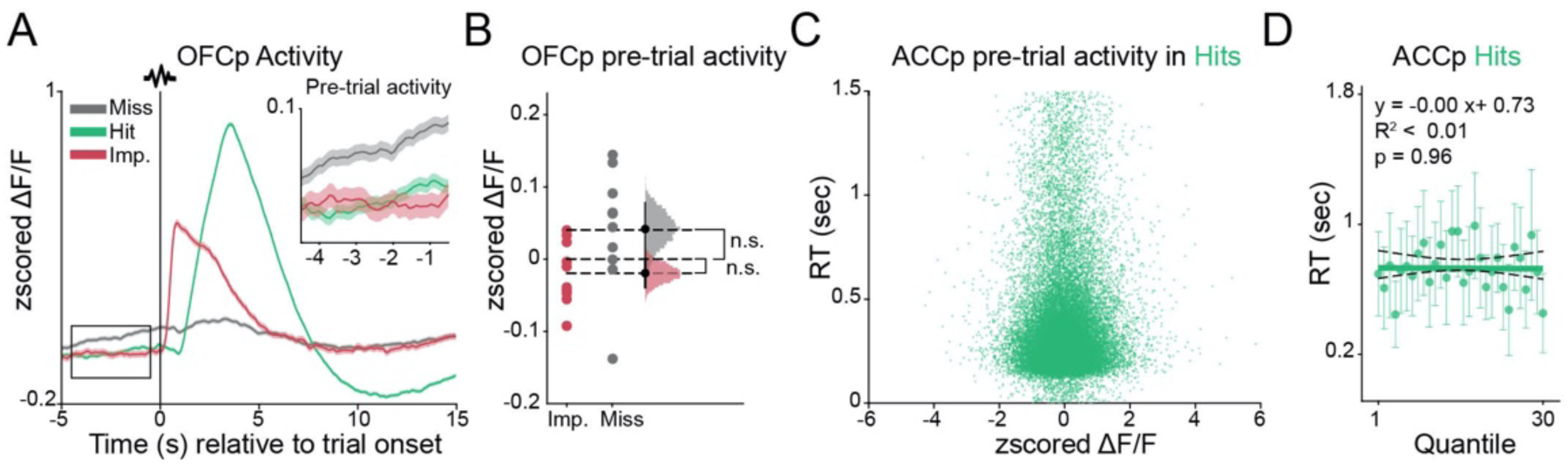
OFCp pre-trial activity is not correlated with performance. (**A**) Mean activity of all OFCp claustrum recordings (n=10) aligned to trial onset, divided by trial outcome. Inset depicts 5 seconds of pre-trial activity. (**B**) OFCp pre-trial baseline activity in impulsive trials (red) and miss trials (gray), depicted as individual mice and bootstrapped distribution of means with 95% confidence intervals (n=10). (**C**) Response time in hits as a function of ACCp pre-trial activity (n=20). (**D**) Response time in ACCp mice (n=20) is uncorrelated with pre-trial activity. Thick line represents linear fit, dotted lines represent 95% confidence intervals. Unless noted otherwise, data are mean ± s.e.m. *p < 0.05, **p < 0.01, ***p < 0.001; n.s., not significant. See Supplementary Table 3 for further details of the statistical analyses.

**Figure S7.**
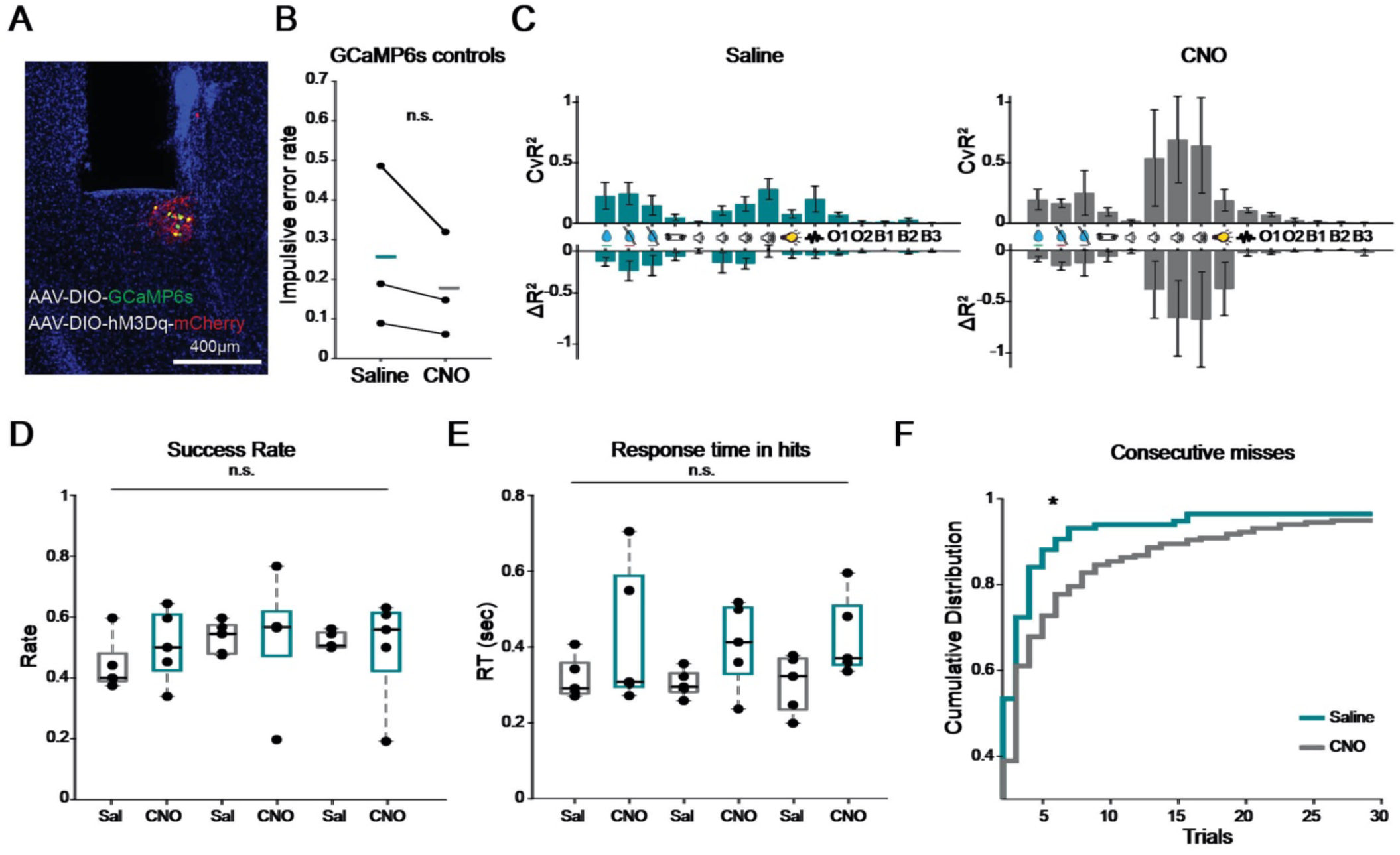
The effect of ACCp chemogenetic activation is specific, and does not affect transient responses to task events, while enhancing consecutive misses late in the session. (**A**) Expression of GcAMP6s (green) and hM3Dq (red) in ACCp neurons. (**B**) CNO had no effect on impulsive errors in GCAMP6s control mice (n=3). (**C**) Model quantification of the contribution of behavioral events to claustrum photometry signals in saline (left) or CNO (right) sessions (n=5 mice, 3 sessions of each condition per mouse). (**D**) Success rates of ACCp-hM3Dq expressing mice throughout the experiment were not affected by CNO. (**E**) Response time in hit trials throughout the experiment were not affected by CNO. Boxes in (**D, E**) represent group median and 1st and 3rd quartiles, session order as in Figure 3D. (**F**) Cumulative probability distribution of consecutive miss trials in saline (turquoise) and CNO (gray) sessions. Unless noted otherwise, data are mean ± s.e.m. *p < 0.05, **p<0.01, ***p< 0.001; n.s., not significant. See Supplementary Table 3 for further details of the statistical analyses.

**Figure S8.**
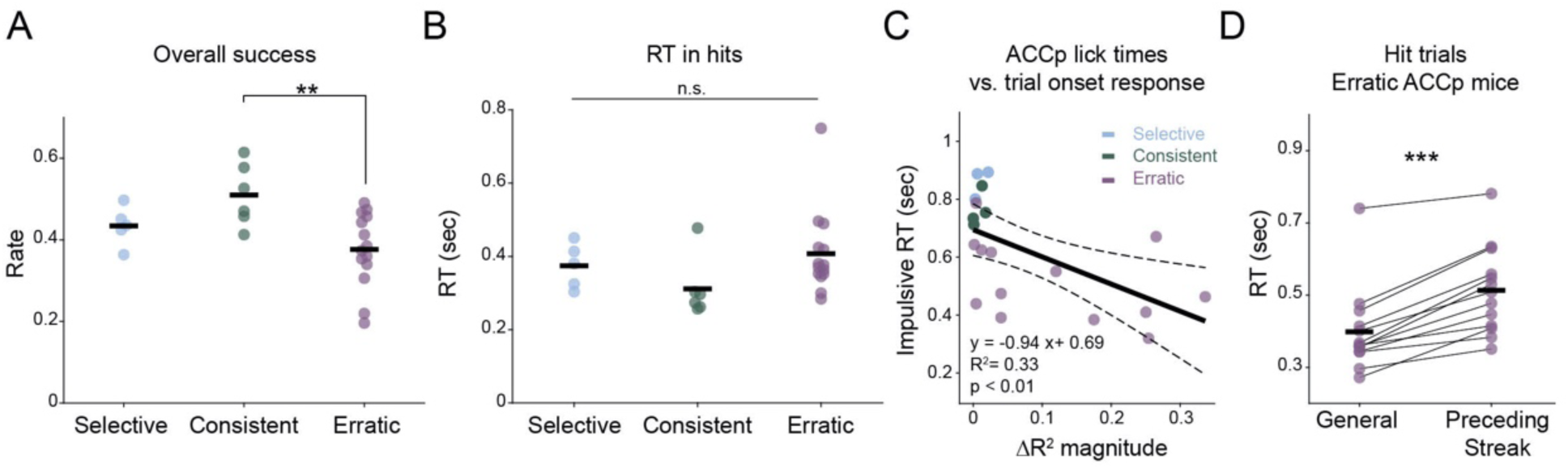
Erratic mice exhibit an overall lower success rate, and increased reaction time prior to miss streaks. (**A**) Overall success rate in the task by strategy group (n = 5 selective; 6 consistent; 14 erratic). (**B**) No relation of reaction time in hit trials to strategy group. (**C**) Mean response time in impulsive errors (Figure 4E) as a function of the model quantification for trial onset ACCp activity (Figure 4F). Colors represent strategy group (n = 3 selective; 4 consistent; 13 erratic). Thick line represents linear fit, dotted lines represent 95% confidence intervals. (**D**) Response time for erratic ACCp mice (n=13) is increased in hit trials immediately preceding miss streaks, compared to all hit trials. Unless noted otherwise, data are mean ± s.e.m. *p<0.05, **p<0.01, ***p< 0.001; n.s., not significant. See Supplementary Table 3 for further details of the statistical analyses.

**Figure S9.**
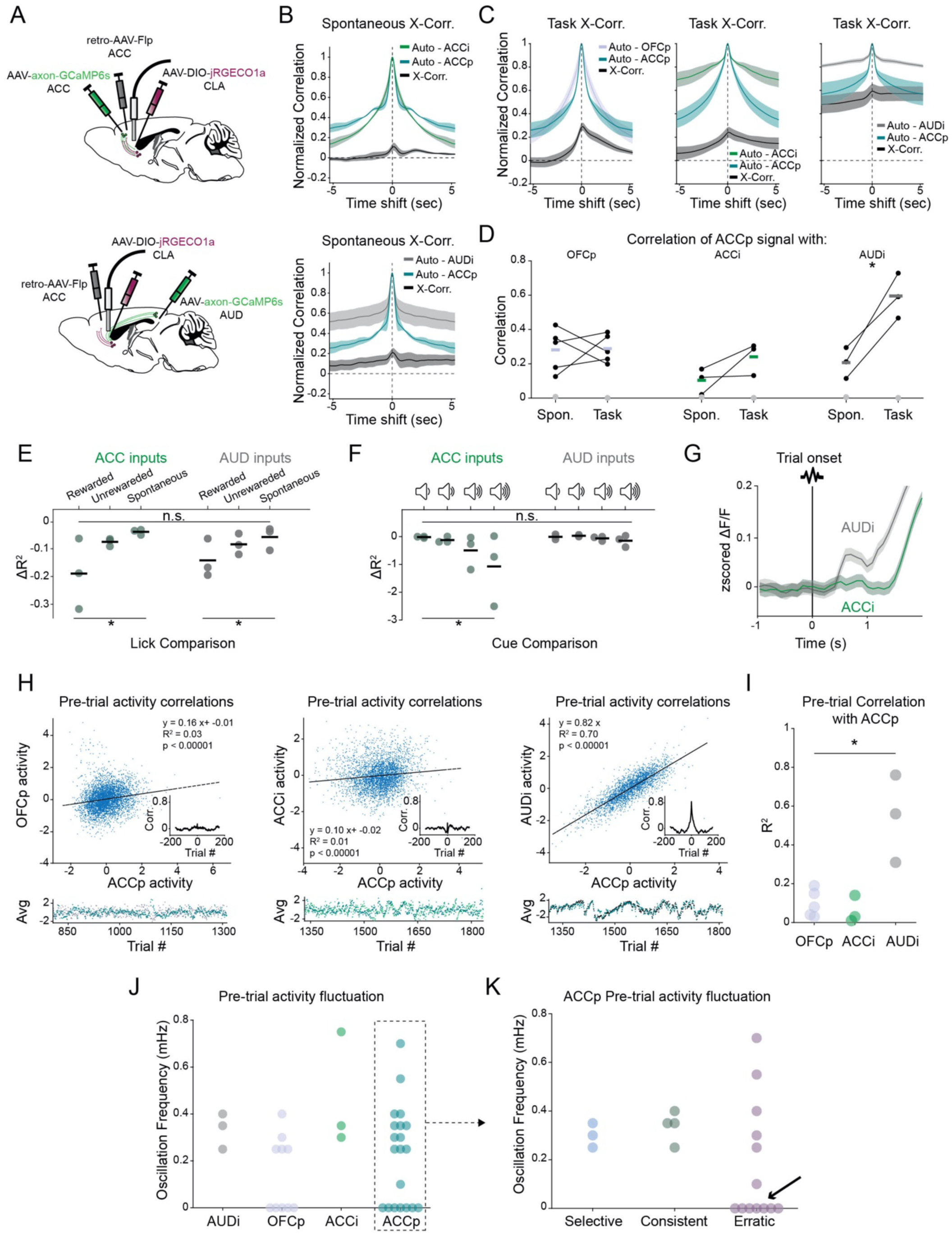
Claustral inputs from auditory cortex acquire task-dependent correlations with ACCp. (**A**) Strategy for simultaneous recording from claustrum ACCp neurons expressing jRGECO1a, together with activity of claustrum afferents from ACC (ACCi; top) or AUD (AUDi; bottom) axons, using axonal-targeted GCaMP6s. (**B**) Average autocorrelations of ACCi/ACCp or AUDi/ACCp spontaneous activity, and respective cross-correlations between channels (n=3 mice in each group). (**C**) Average autocorrelations of ACCp/OFCp (left), ACCi/ACCp (middle) or AUDi/ACCp (right) activity during task recordings, as well as cross-correlations between channels (n = 5, 3, 3 mice, respectively). (**D**) Summary of overall correlations between channel activity during free recordings (spontaneous) compared to those recorded during the task (n=5, 3, 3 mice for each group), demonstrating an increase in correlated activity between AUDi and ACCp networks during the task. Gray dots represent the maximal correlation of shuffled data over 1000 iterations per mouse, averaged across mice. (**E-G**) Model quantification of ACC (ACCi) and auditory (AUDi) cortical inputs to the claustrum. (**E-F**) Quantification of the contribution of activity during licking events (**E**) and go cue stimuli (**F**) to the overall signal. (**G**) Average trace from all axonal recordings in ACCi (green) vs AUDi (gray), aligned to trial onset, depicting trial onset responses in AUDi, and their absence in ACCi activity. (**H-K**) Pretrial activity dynamics. (**H**) Correlation of average pre-trial activity (5s preceding trial onset) in representative co-recorded ACCp/OFCp (left), ACCp/ACCi (middle), or ACCp/AUDi (right) channels. Inset depicts cross-correlation in a window spanning 400 trials. Bottom panel depicts the magnitudes of pre-trial ACCp activity and corresponding OFCp, ACCi or AUDi activity during individual consecutive trials in a representative session. (**I**) Coefficient of determination (R-squared) of the linear fit between pre-trial activity (5 s prior to trial onset) in co-recorded channels. (**J**) Frequency of ultraslow oscillations of pre-trial activity in ACCp (n=20), OFCp (n=10), ACCi (n=3) and AUDi (n=3) recordings. Oscillation frequency was defined as the peak of the frequency spectrum emerging above a threshold obtained from 1000 shuffles of the data (see Methods). (**K**) Division of pre-trial ultra-slow fluctuations in ACCp mice according to strategy (see **Figure 4**). Arrow points to mice with no significant oscillation, all associated with the erratic group. Unless noted otherwise, data are mean ± s.e.m. *p < 0.05, **p < 0.01, ***p < 0.001; n.s., not significant. See Supplementary Table 3 for further details of the statistical analyses.

**Figure S10.**
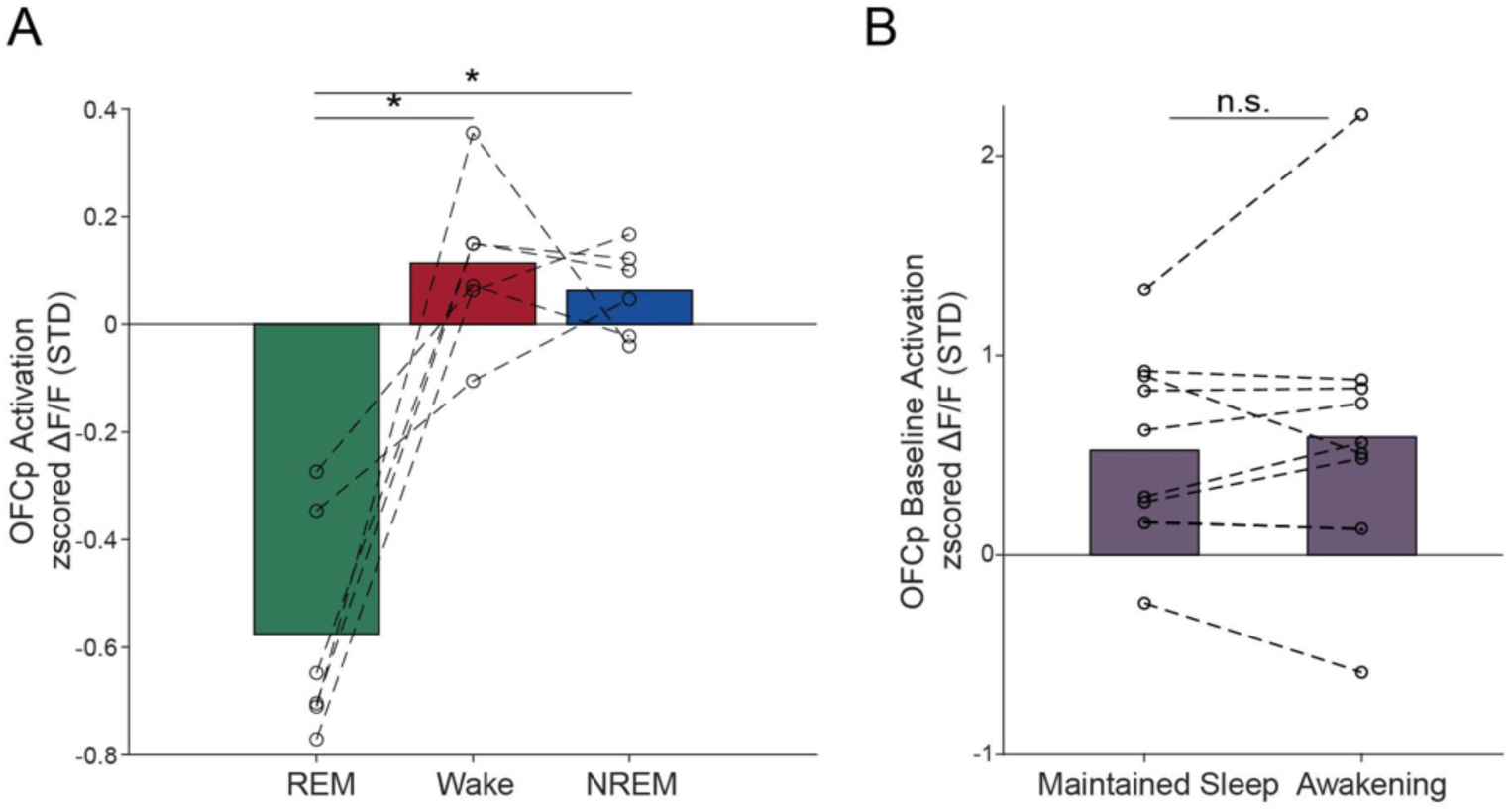
Claustral OFCp activity during sleep. (**A**) Average OFCp claustrum calcium activity in REM sleep, wake, and NREM sleep (n=6). Note that in this population, activity during NREM sleep is not higher than in wakefulness. (**B**) OFCp baseline activity (y-axis) for trials associated with maintained sleep (left) vs. awakening (right). Each dot represents a separate ∼10h experiment (10 experiments in n=6 mice). Unless noted otherwise, data are mean ± s.e.m. *p < 0.05, **p < 0.01, ***p < 0.001; n.s., not significant. See Supplementary Table 3 for further details of the statistical analyses.

**Supplementary Table T3.**
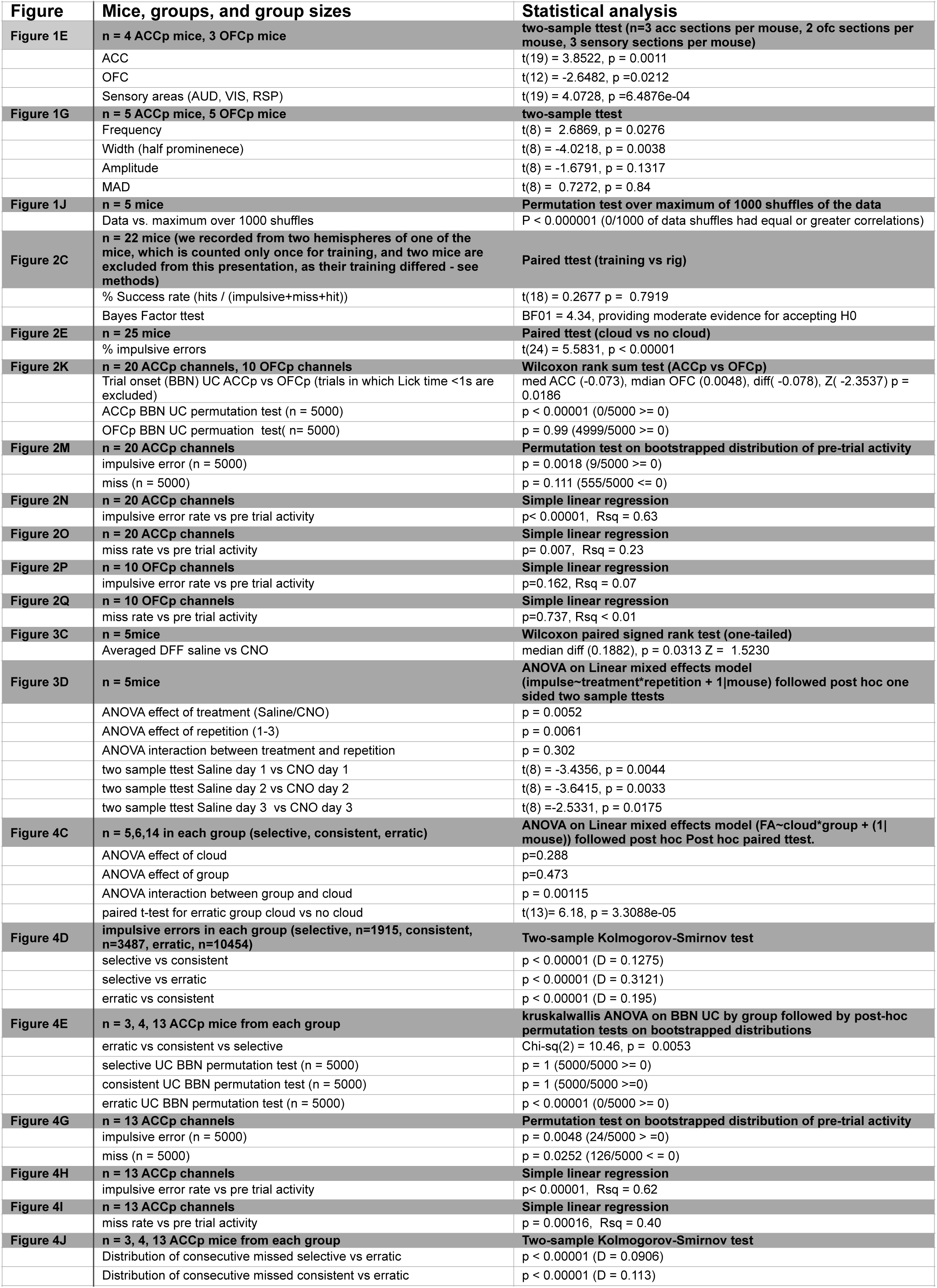
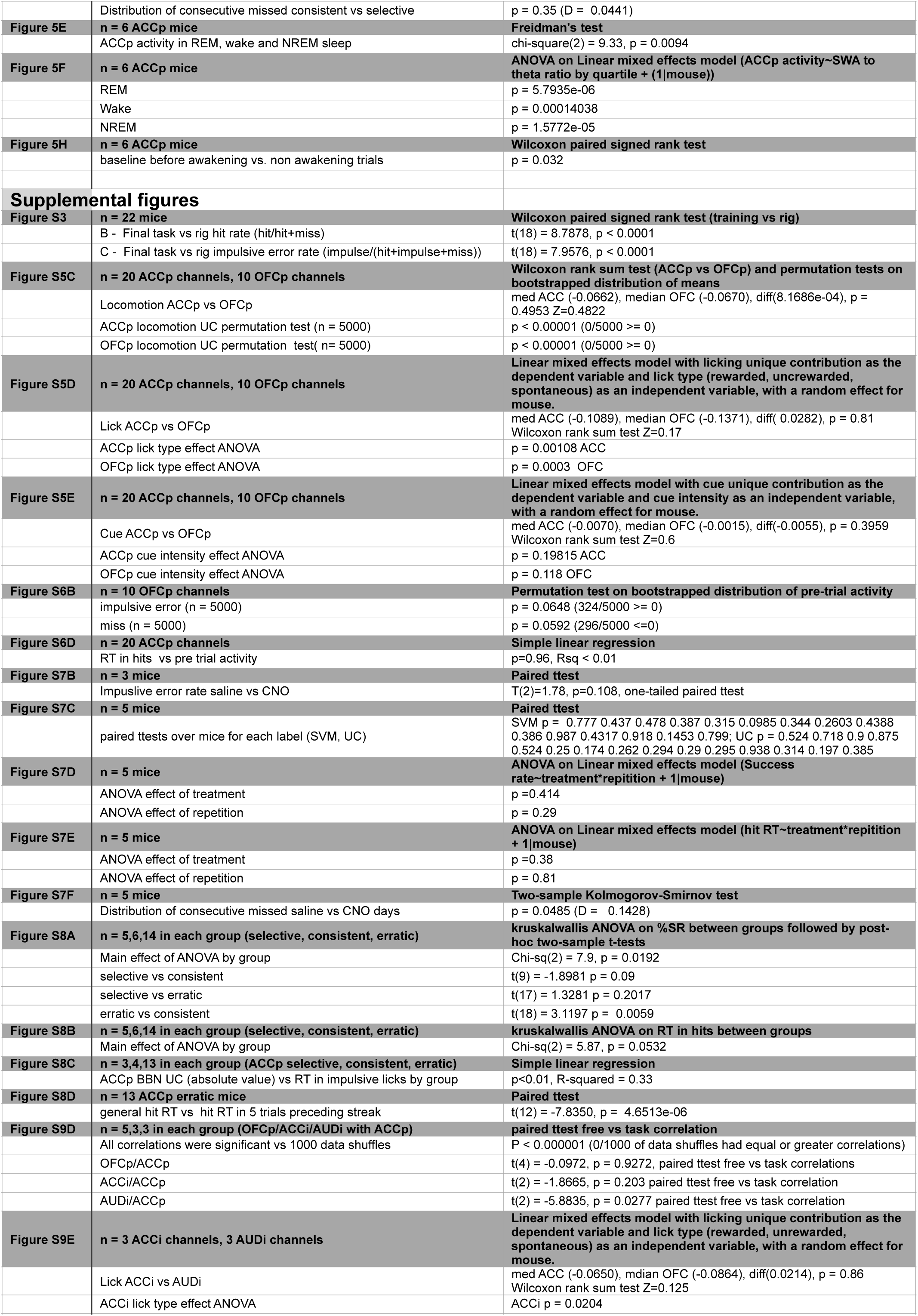
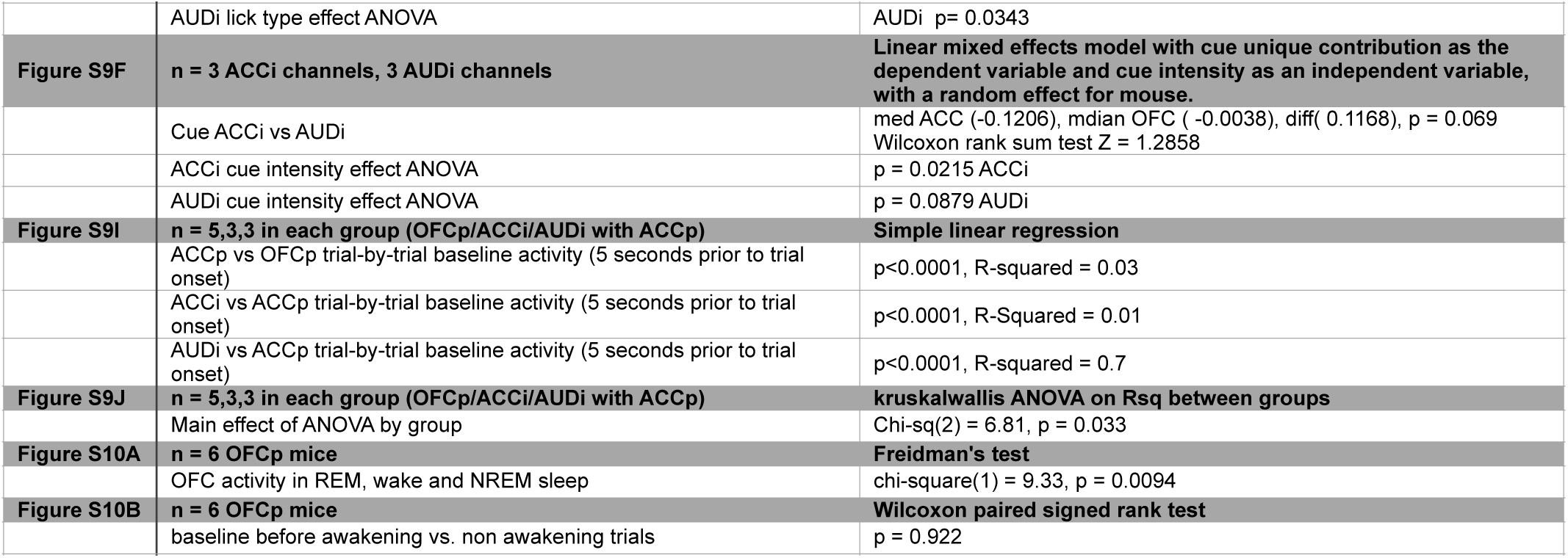
Statistical Analysis

